# Targeting the cell and non-cell autonomous regulation of 47S synthesis by GCN2 in colon cancer

**DOI:** 10.1101/2023.02.08.527626

**Authors:** Marie Piecyk, Mouna Triki, Pierre-Alexandre Laval, Cedric Duret, Joelle Fauvre, Laura Cussonneau, Christelle Machon, Jerôme Guitton, Nicolas Rama, Benjamin Gibert, Gabriel Ichim, Frederic Catez, Fleur Bourdelais, Sebastien Durand, Jean-Jacques Diaz, Isabelle Coste, Toufic Renno, Serge N Manié, Nicolas Aznar, Stephane Ansieau, Carole Ferraro-Peyret, Cedric Chaveroux

**Affiliations:** Centre de Recherche en Cancérologie de Lyon, INSERM U1052, CNRS 5286, Centre Léon Bérard, Université de Lyon, Université Claude Bernard Lyon 1, Lyon, France; Université Clermont Auvergne, INRAE, Unité de Nutrition Humaine, UMR1019, Clermont-Ferrand, France; Biochemistry and Pharmaco-toxicology laboratory, Lyon Sud Hospital, University Hospital of Lyon, Hospices Civils de Lyon Pierre-Bénite, France; Hospices Civils de Lyon, Plateforme AURAGEN, Lyon, France

## Abstract

Nutrient availability is a key determinant of tumor cell behavior. While nutrient-rich conditions favor proliferation and tumor growth, scarcity, and particularly glutamine starvation, promotes cell dedifferentiation and chemoresistance. Here, linking ribosome biogenesis plasticity with tumor cell fate, we uncover that the amino acid sensor GCN2 represses the expression of the precursor of ribosomal RNA, 47S, under metabolic stress. We show that blockade of GCN2 triggers cell death by an irremediable nucleolar stress and subsequent TP53-mediated apoptosis in patient-derived models of colon adenocarcinoma (COAD). In nutrient-rich conditions, GCN2 activity supports cell proliferation through the transcription stimulation of 47S rRNA, independently of the canonical ISR axis. However, impairment of GCN2 activity prevents nuclear translocation of the methionyl tRNA synthetase (MetRS) underlying the generation of a nucleolar stress, mTORC1 inhibition and autophagy induction. Inhibition of the GCN2-MetRS axis drastically improves the cytotoxicity of RNA pol I inhibitors, including the first-line chemotherapy oxaliplatin, on patient-derived COAD tumoroids. Our data thus reveal that GCN2 differentially controls the ribosome biogenesis according the nutritional context. Furthermore, pharmacological co-inhibition of the two GCN2 branches and the RNA pol I activity may represent a valuable strategy for elimination of proliferative and metabolically-stressed COAD cell.

## Introduction

Cancer cell proliferation implicates a rewired metabolism to sustain the generation of biomass (Pavlova and Thompson, 2016). This cell growth is concomitantly supported by enhanced ribosome biogenesis (RiBi). However, in solid tumors, variation of nutrient availability determines protein homeostasis and the balance between proliferation capacity or quiescence. Hence, nutrient availability is often found in lower concentrations in various tumors, including colon adenocarcinoma (COAD), compared to the normal tissue or in tumor interstitial fluid compared to the plasma (Kamphorst et al., 2015; Sullivan et al., 2019). Fluctuations in amino acids availability also occur depending on the tumor region (Pan et al., 2016). Shifting cell fate from a proliferative to a quiescent state thus relies on the proteostatic flexibility, notably implicating a fine tuning of the RiBi. Nonetheless, the sensors and downstream mechanisms orchestrating the adaptation of RiBi according the nutritional context remain unclear.

RiBi depends on the production of ribosomal proteins (riboproteins) encoded by RNA polymerase II (pol II) and of the 5S ribosomal RNA (rRNA) encoded by RNA pol III. In addition, RNA pol I transcribes the 47S precursor rRNA (pre-rRNA), which will then be processed into final rRNA products, namely the 5.8S, 18S and 28S (Catez et al., 2019). The latter mechanism, occurring within the nucleolar sub-compartment, is sensitive to nutritional variations and is disrupted upon starvation, making the nucleolus a stress-integrator site (Clarke et al., 1996). Ultimately, defects in RiBi trigger nucleolar stress leading to cell cycle arrest and/or apoptosis (Russo and Russo, 2017).

Adjustment of the translation rate according the nutritional milieu implicates a tight reciprocal regulation between the ribosome biogenesis and nutrient sensor such as the mammalian target of rapamycin complex 1 (mTORC1). Active mTORC1 is commonly found in many types of cancers (Edinger and Thompson, 2002; Harachi et al., 2018; Saxton and Sabatini, 2017). In nutrient rich conditions, this complex supports protein synthesis and tumor progression notably by enabling RNA pol I and III transcription, rRNA processing and riboprotein translation (Guo et al., 2018; Sameer, 2013). However, in poorly-perfused areas, mTORC1 inactivation by the lack of nutrients contributes to RiBi repression. (Gomes et al., 2016; Riedl et al., 2017; Riffle and Hegde, 2017). Interestingly, the RNA pol I activity reciprocally controls the mTORC1 pathway. Indeed, decreased RNA pol I activity and impairment of 47S synthesis is sufficient to downregulate mTORC1 that in turns induces autophagy (Chen et al., 2016; Goudarzi et al., 2014; Li et al., 2016; Zajkowicz et al., 2015). Inactive mTORC1 orchestrates a translational reprogramming alongside the phosphorylation of the eukaryotic initiation factor 2a (P-eIF2a) on serine 51 (Jewer *et al*, 2020). This latter triggers the integrated stress response (ISR) and has two major consequences: i) a repression of translation initiation and, ii) up-regulation of the translational of specific transcripts embedding open reading frames (uORFs) in their 5’ untranslated region, including the activating transcription factor 4 (ATF4). ATF4 then orchestrates a transcriptional program to restore amino acids homeostasis, angiogenesis and antioxidant response to ultimately promote cell survival (Chaveroux et al., 2016; Harding et al., 2003; Sarcinelli et al., 2020; Wang et al., 2013). Following nutrient scarcity or proteotoxic stresses, GCN2 is the only eIF2a kinase activated by ribosome collision (Dong et al., 2000; Harding et al., 2019; Inglis et al., 2019). Once activated, GCN2 participates to the adaptive translational and metabolic response by canonically inducing the ISR. However the contribution of GCN2 in the regulation of the ribosome biogenesis is poorly described. Cells silenced for GCN2 display an elevation of the RNA pol III activity and notably 5S rRNA (Nakamura and Kimura, 2017). Yet accumulation of 5S, in complex with RPL5 and RPL11, occurs subsequently to an impairment of the generation of 47S and related sub-products, suggesting that augmentation of the 5S content in GCN2-knockdown likely arise from a prior dysregulation of the 47S production and/or processing (Sloan et al., 2013). Furthermore, the impact of the nutritional milieu on the GCN2-RiBi interplay was still unclear. Indeed, previous work by our group and others intriguingly suggested that GCN2 may also regulate cell viability and migration even when nutrients are abundant through an unknown mechanism (Chaveroux et al., 2016; Ge et al., 2018; Li et al., 2022; Nakamura and Kimura, 2017).

Our present study is based on the hypothesis that GCN2 may differentially regulate the rate of protein synthesis in cancer cells through the modulation of 47S metabolism in quiescent and proliferative cells. Taking COAD as a paradigm of cancers displaying ribosomal dysfunctions associated to amino acid starvation, we show that, in a harsh nutritional context, GCN2 actively contributes to COAD cancer cell plasticity through the repression of 47S pre-rRNA synthesis, constituting a barrier against an irreversible nucleolar stress and ultimately cell death. Furthermore, in nutrient-abundant condition, we show that inhibition of GCN2 prevents the nuclear translocation of the methionyl tRNA synthetase (MetRS) and impairs the 47S pre-RNA expression, further associated to mTORC1 repression and autophagy induction. This ribosomal vulnerability caused by GCN2 inhibition confers a higher sensitivity to RiBi stressors including the chemotherapeutic agent oxaliplatin. Our results unveil distinct cell-autonomous and non-autonomous functions for GCN2 according to the nutritional milieu both concurring to the maintenance of RiBi and COAD progression.

## Results

### I. Dysregulation of ribosomal function is associated with amino acid starvation and ATF4 activity in COAD

To investigate the crosstalk between the altered nutritional microenvironment, activated GCN2 pathway and ribosomal stress, we first analyzed cancers that displayed clear proteostasis and/or ribosomal impairment. Comparison of normal and tumor samples from several cohorts in the TCGA database (figure supplement 1A) identified COAD as having highly dysregulated ribosome biogenesis (RiBi) and cell proliferation activities (Figure 1A). Defects in ribosome assembly in human tumors arise following impairment of rRNA production, or riboprotein (RP) processing and assembly (Albert et al., 2019; Kim et al., 2014). We then compiled a gene signature of RPs with a significantly dysregulated expression (negatively and positively) and confirmed enrichment of this signature in COAD compared to the normal tissue (table supplement 1) (Guimaraes and Zavolan, 2016). As expected, this signature was, validating its relevance for assessing ribosomal stress in this model (Figure 1B). To determine whether RP dysregulation was associated with metabolic stress, we compared enrichment of previously described signatures related to hypoxia or amino acid starvation in COAD based on two levels of RP dysregulation, low or high (Figure 1C, table supplement 1) (Dekervel et al., 2014; Krige et al., 2008). Interestingly, patients with a high RP dysregulation also displayed a significant enrichment in the amino acid starvation signature, albeit no difference was observed for the COAD-specific hypoxia signature between the two groups (figure supplement 1B), indicating that ribosomal stress may be strongly associated with a lack of amino acids. At the molecular level, amino acid deprivation and ribosomal stress are both strong inducers of the GCN2 kinase triggering the ISR-driven transcriptional program. We then assessed in the same subgroups of patients, a gene signature related to ATF4 transcriptional activity, the terminal factor of the GCN2-ISR pathway (table supplement 1). We first confirmed that the ATF4 signature was enriched in tumors compared to the normal tissue (Figure 1D). A correlation analysis then confirmed that this signature was linked to the amino acid deprivation signature (figure supplement 1C), and was higher in COAD tumors with a high RP dysregulation (Figure 1E). The data thus identified ATF4 as a major contributor to the stress response related to amino acid deprivation in COAD patients.

**Fig. 1.**
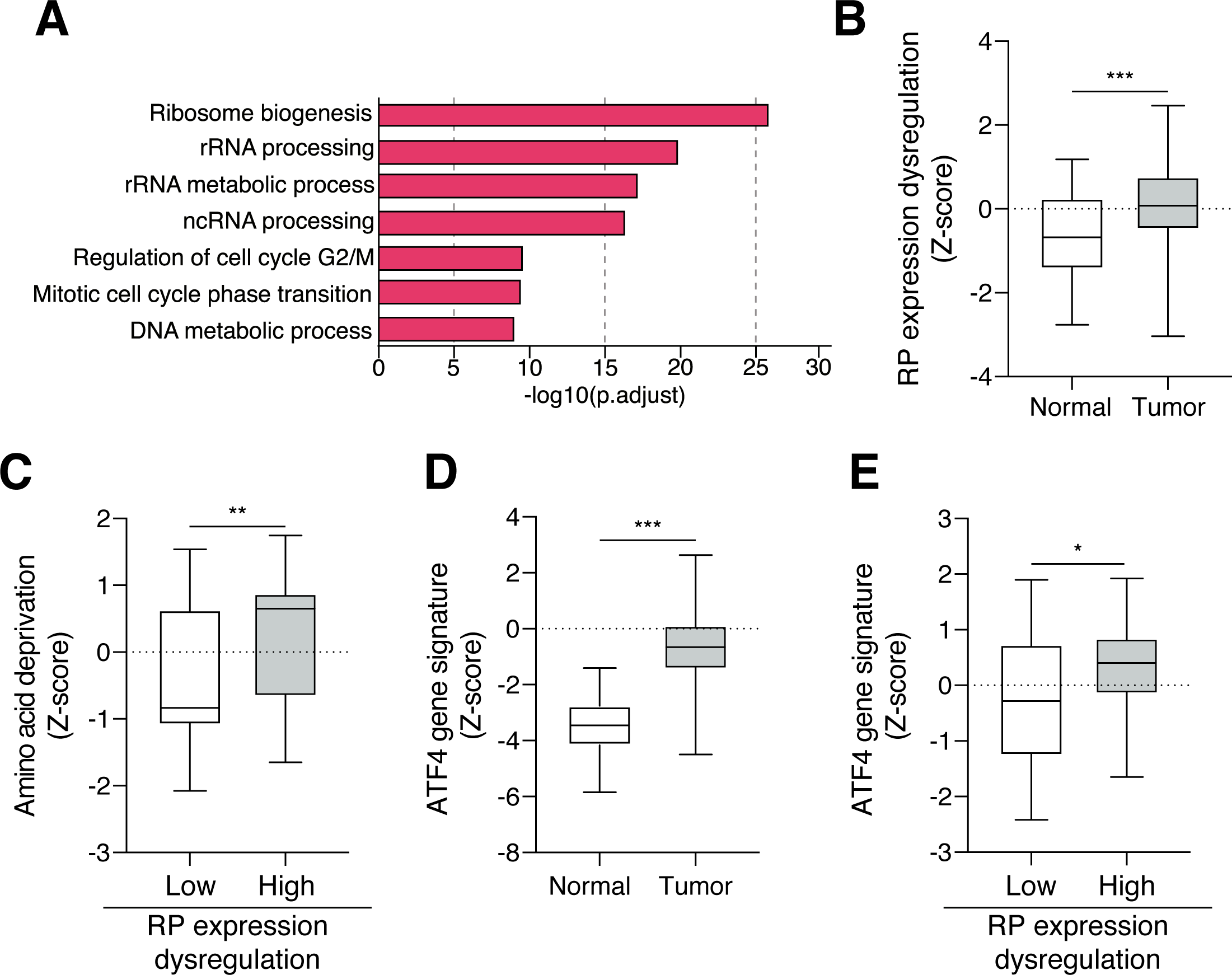
Ribosomal dysfunction in colon adenocarcinoma (COAD) is correlated with the upregulation of ATF4 activity. A. Gene Ontology enrichment analysis on TCGA database highlights an upregulation of ribosomal biogenesis processes in COAD compared to normal tissue. Top enrichment GO terms in COAD tumors versus normal tissues are shown and significance is expressed as -Log10 adjusted p-value. B. Dysregulated expression of riboproteins (RP) is higher in COAD compared to the normal tissue. TCGA COAD data, Mann-Whitney-test, (*** p < 0.001) C. The amino acid deprivation gene signature is enriched in COAD tumors with high dysregulated RP expression. TCGA COAD data, Mann-Whitney-test, (** p < 0.01) D. ATF4 gene signature is higher in colon tumors compared to normal tissue. TCGA COAD data, Mann-Whitney-test, (*** p < 0.001) E. ATF4 gene signature is enriched in COAD with high dysregulation of RP expression. TCGA COAD data, Mann-Whitney-test (* p < 0.05)

### II. GCN2 represses 47S pre-RNA expression upon glutamine scarcity and prevents TP53-dependent apoptosis

GCN2 takes part in survival upon ribotoxic stress. Here, we sought to decipher its role in maintaining cell viability and nucleolar integrity in those harsh conditions. We first confirmed the accumulation of GCN2 at the mRNA and protein levels, in a large group of patients and even at early stages of the pathology (figure supplement 2A-C, Figure 2A, B) (Marisa et al., 2013). Clinically, high levels of GCN2 expression in COAD patients were correlated with a lower disease-specific survival (figure supplement 2D). Considering the role of the ISR in cancer cell survival through cell plasticity in a nutrient-deprived microenvironment, we then set up *in vitro* conditions to assess whether GCN2 impacted COAD cell plasticity. HCT116 and LoVo COAD cells were grown as three dimensional (3D) models in order to mimic the complex metabolic heterogeneity and physiochemical gradients of solid tumors (Nath and Devi, 2016). Spheroids were then treated with a previously published GCN2 inhibitor, TAP20, hereafter referred to as GCN2i (Bröer et al., 2019). Prior validation in 2D culture confirmed the ability of GCN2i to abrogate the accumulation of ATF4 and related target genes (*CHOP, TRB3, GADD34, ASNS, SESN2*) upon leucine and glutamine (Gln) starvation (figure supplement 3A-C). In spheroids, viability assays following GCN2i treatment highlighted the presence of dead cells in the central region of spheroids confirming the protective role of GCN2 against an altered metabolic microenvironment and its implication in cell plasticity (Figure 2C, D). COAD patient-derived tumoroids were then grown as colospheres to reproduce the metabolic gradients (figure supplement 4) and our results were also confirmed by visualizing cell death in the compact core of the 3D models (Figure 2E). In contrast, the addition of the Rho-associated kinase (ROCK) inhibitor (Y-27632), a compound commonly present in tumoroids culture medium to inhibit cell death and enhance stem cell growth, abrogates the induction of cell death by the GCN2i (Kan et al., 2015; Ohata et al., 2012). Immunohistological analyses further confirmed the loss of cell viability and enhancement of apoptosis upon GCN2 inhibition in spheroids core, as attested by the measurement of cleaved PARP and cleaved caspase-3 staining (Figure 2F, G). Finally, the expression of two COAD stemness-related markers, *ALDH1A3* and c-*MYC* (Elbadawy et al., 2019; Gelardi et al., 2021), was reduced in spheroids treated with GCN2i, confirming that GCN2 is essential for the viability of c-Myc-expressing tumor cells and the elimination of quiescent tumor cells in the presence of the drug (Figure 2H).

**Fig. 2.**
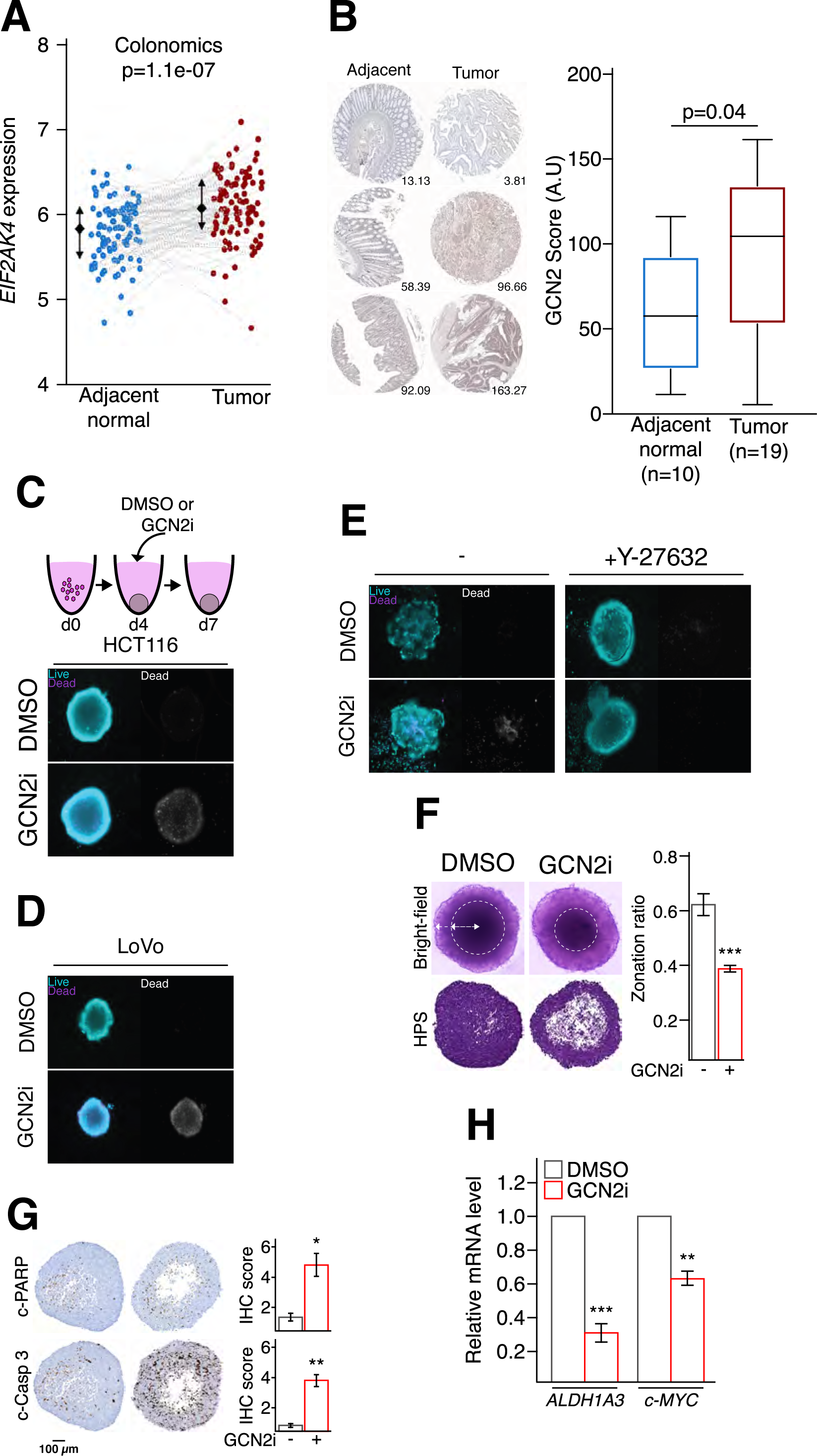
GCN2 activity is critical to promote COAD cells plasticity. A. *GCN2* (gene name *EIF2AK4*) expression levels is higher in colon tumor tissues compared to the normal counterparts. Colonomics cohort, Unpaired t-test. B. Immunohistochemistry analysis confirmed that GCN2 amount is augmented in human COAD tumors compared to adjacent normal tissues. Quantification is provided on the right panel. Unpaired t-test. C. Live-and-dead assay in HCT116 spheroids treated for 3 days with vehicle (DMSO) or GCN2i. To facilitate the observation of death induction, staining of non-viable cells is also represented in shades of grey (right panel). D. Live-and-dead assay in LoVo spheroids treated for 3 days with vehicle (DMSO) or GCN2i. To facilitate the observation of death induction, staining of non-viable cells is also represented in shades of grey (right panel). E. Live-and-dead assay in patient’s derived colospheres, grown with or without the ROCK inhibitor (Y-27632), after 3 days of GCN2i treatment. To facilitate the visualization of cell death induction, staining of non-viable cells is also represented in shades of grey (right panel). F. Evaluation of cellular compaction in the core of HCT116 spheroids treated for 3 days with DMSO or GCN2i via bright-field microscopy and on hematoxylin-phloxine-saffron stained spheroid slices. Data are expressed as mean +/-s.e.m of independent experiments (n = 3), unpaired two-tailed t-test with the p-value (*** p < 0.001). G. Measurements of immunohistochemical staining against cleaved PARP and cleaved caspase 3 on HCT116 spheroids treated with DMSO or with GCN2i for 3 days. Data are expressed as mean +/-s.e.m of independent experiments (n = 3), unpaired two-tailed t-test with the p-value (* p < 0.05, ** p < 0.01). H. RT-qPCR analysis of *ALDH1A3* and *c-MYC* mRNA levels in HCT116 spheroids treated for 48h with DMSO or GCN2i. Data are expressed as mean +/-s.e.m of independent experiments (n = 3), unpaired two-tailed t-test with the p-value (** p < 0.01, *** p < 0.001).

Having validated that COAD represent a relevant context for studying the crosstalk between the GCN2 pathway and the RiBi. Then we determined whether this kinase is implicated in maintaining the nucleolar integrity under low amino acid conditions. Mechanistical investigations required to shift toward simpler cellular models. In 2D culture, cells were exposed to low concentrations of asparagine or glutamine (Gln) similar to those found in poorly-perfused area of solid tumors. Cell death measurement revealed that GCN2i drastically killed the cells once associated to glutamine but not asparagine, (figure supplement 5A) (Pan et al., 2016). Combination with inhibitors of necroptosis (R3i, Nec1) or apoptosis (QVD, Z-VAD) confirmed that the loss of viability was caused by apoptosis when Gln starvation and GCN2i were combined (figure supplement 5B). Since the results obtained in 2D mirrored our observations in 3D, we hereafter applied the same Gln-deprived conditions to evaluate whether nucleolar integrity was impaired. A redistribution of fibrillarin, a canonical nucleolar marker, was observed in starved cells upon GCN2i indicating the induction of a nucleolar stress (Figure 3A) (Yuan et al., 2005). To document if this stress is a consequence of an impairment of the RNA pol I activity, northern blots were performed to assess 47S synthesis (Figure 3B). Gln-starved cells displayed a lower amount of 47S compared to the control, which is consistent with an inhibition of Pol I activity upon stress (Grummt et al., 1976). However, in presence of GCN2i, the level of 47S pre-rRNA increased again indicating the recovery of RNA pol I activity. To assess whether cell death caused by GCN2 inhibition upon low glutamine is a consequence of the nucleolar stress, the amount of TP53 was monitored in the same conditions. Indeed, TP53 is widely described as the molecular effector between nucleolar stress caused by RNA pol I repression and apoptosis (Fumagalli et al., 2012; James et al., 2014; Rubbi and Milner, 2003; Yang et al., 2018). Consistent with previous study, low glutamine alone did not change the level of TP53 during the starvation time (Lowman et al., 2019). However, cotreatment with GCN2i triggered an elevation of TP53 amounts and its target gene MDM2 (Figure 3C), two canonical markers of nucleolar stress (Morgado-Palacin et al., 2012). To functionally confirm whether GCN2i-induced cell death relies on TP53 upon, live-and-dead analyses were performed in spheroids made from COAD cell lines expressing a wild type (HCT116, LoVo) or mutant (HT29 and DLD-1) forms of TP53 (figure supplement 5C). Although a loss of cancer cell viability was obviously observed in the core area of LoVo and HCT116 spheroids upon GCN2i, the treatment did not lead to cell death in the spheroids mutated for TP53. To avoid a comparison bias related to the heterogenous capacities of the cell lines to form comparable spheroids, we shifted back to 2D cultures (figure supplement 5D). GCN2i treatment diminished cell viability of the TP53 WT cells upon low Gln whereas TP53 mutant cells were not affected by the blockade of GCN2 activity. Furthermore, toxicity assays performed on the isogenic HCT116 cell lines WT or knockout (KO) for TP53 confirmed, in 3D and 2D, that this latter is required for cell death induction by the drug upon deprivation (Figure 3D, E). Of note, absence of TP53 did not impact the induction of ATF4 upon low Gln (figure supplement 5E). Altogether, these results demonstrate that GCN2 critically contributes to the RiBi repression upon amino acids scarcity thus preventing a TP53-mediated apoptosis.

**Fig. 3.**
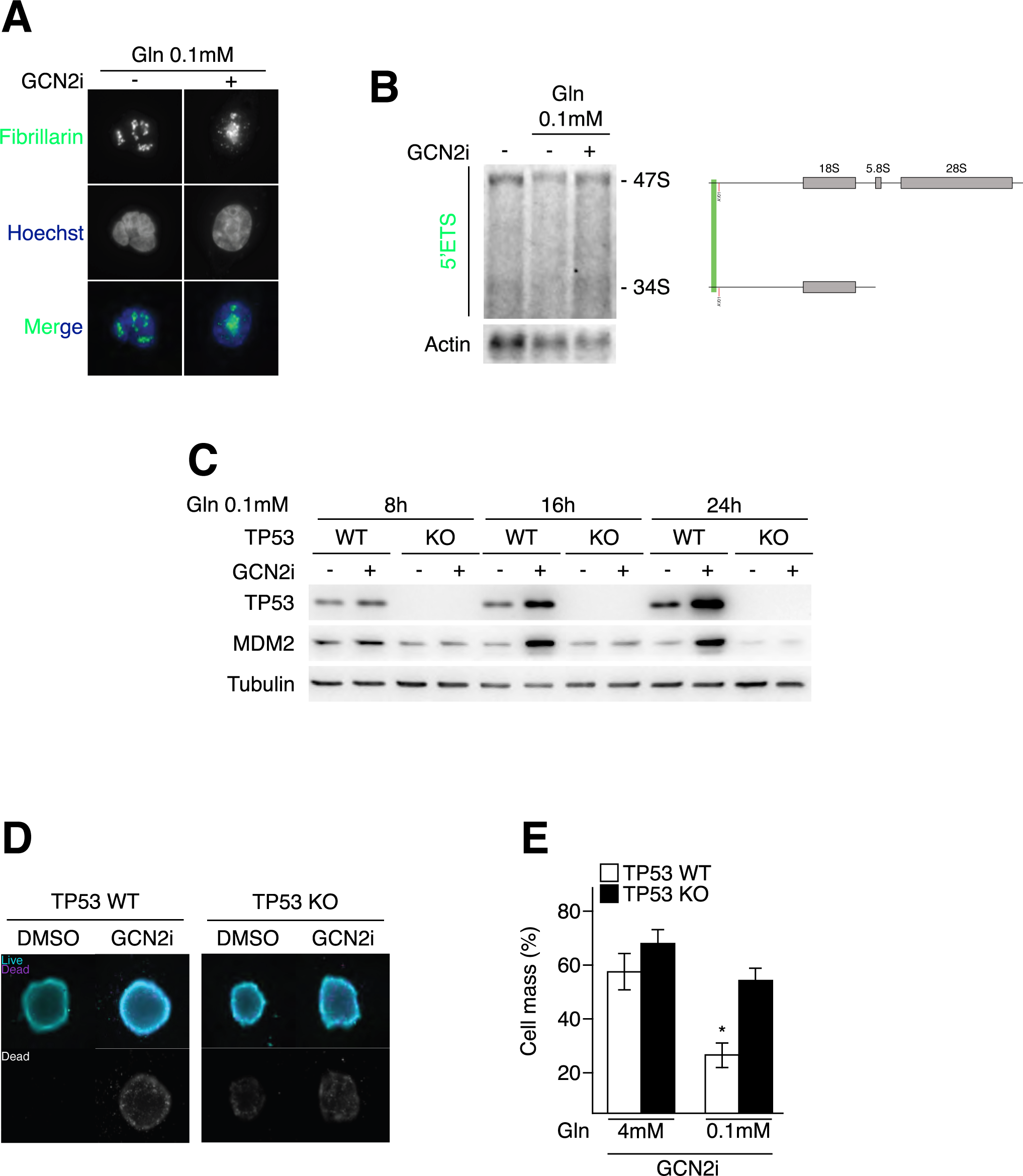
GCN2 is required for repressing 47S transcription and preventing the TP53-proapoptotic program upon low glutamine. A. Fibrillarin localization assessed by immunofluorescence in glutamine-starved HCT116 cells treated with DMSO or GCN2i. B. Northern blotting of cellular rRNAs after indicated treatments. The position of the investigated probes on the 47S rRNA and sub-products are indicated according the corresponding colors. C. Western blot analysis of TP53 and MDM2 protein amounts in HCT116 TP53 WT and HCT116 TP53 KO treated with DMSO or GCN2i upon low glutamine condition (0.1 mM, -Gln) for the indicated period of time. D. Live-and-dead assay in HCT116 spheroids wild-type (WT) or knockout (KO) for TP53 treated for 3 days with vehicle (DMSO) or GCN2i. To facilitate the observation of death induction, staining of non-viable cells is also represented in shades of grey (right panel). E. Measurements of cell mass impairment in HCT116 TP53 WT and HCT116 TP53 KO treated with GCN2i for 48 h in medium containing 4 mM or 0.1 mM levels of glutamine. Data are expressed as mean +/-s.e.m of independent experiments (n = 5). Results of unpaired two-tailed t-test are indicated with the p-value (* p < 0.05).

### III. GCN2, but not the ISR, sustains the mTORC1 signaling in proliferative condition

Having shown the critical role of GCN2 in the maintenance of the RiBi pathway of COAD cells upon nutritional stress, we next investigated its role in nutrient-rich conditions. Indeed, data presented in figure 3E showed that GCN2 inhibition led to a decrease of HCT116 cell mass in nutrient-rich conditions, suggesting that GCN2 may support cell proliferation. To test this, we ibhibited GCN2 in several cell lines, including HCT116, HT29, LoVo and DLD-1 grown under nutrient-balanced conditions (Figure 4A, figure supplement 6A). GCN2i treatment significantly reduced proliferation of HCT116, HT29 and LoVo cells, whereas DLD-1 cells were not affected. To ensure that this impaired proliferation was not due to medium exhaustion, similar experiments were conducted by refreshing the culture medium every day (Figure 4B). Western blot and qPCR analyses confirmed that cells grown in nutrient-rich conditions did not induce the ISR within 24 hours of culture (figure supplement 6B-D). Nonetheless, even in these controlled experimental conditions, we conserved a significant reduction of cell proliferation from 48 hours after GCN2i addition (Figure 4B). This reduction was further confirmed using an RNA interference approach and, pharmacologically, on patient primary COAD cells (Figure 4C, D). Intriguingly, treatment with a second GCN2 inhibitor, GCN2iB, did not impair cell proliferation, suggesting that the two compounds are distinctly modulating GCN2 activity (figure supplement 6E) (Nakamura et al., 2018). Furthermore, GCN2i also impaired spheroids formation indicating that proliferation promotion by this kinase is not restricted only to models in 2D (Figure 4E, figure supplement 6F). To evaluate if GCN2 supports cell proliferation in an ISR-dependent manner, HCT116 cells were treated with ISRIB, a chemical leading to the blockade of eIF2a phosphorylation, or transfected with a siRNA against ATF4 (Figure 4F-I) (Rabouw et al., 2019; Sidrauski et al., 2015). As expected, both treatments prevented the P-eIF2a-dependent elevation of ATF4 and its target gene CHOP in cells deprived for leucine, confirming that the pathway was impaired by the treatment. However, neither ISRIB treatment nor knocking-down ATF4 compromised cell proliferation after 48 hours of treatment. Altogether, these data demonstrate that GCN2 promotes cell proliferation independently of its canonical pathway in abundant-nutrient condition.

**Fig. 4.**
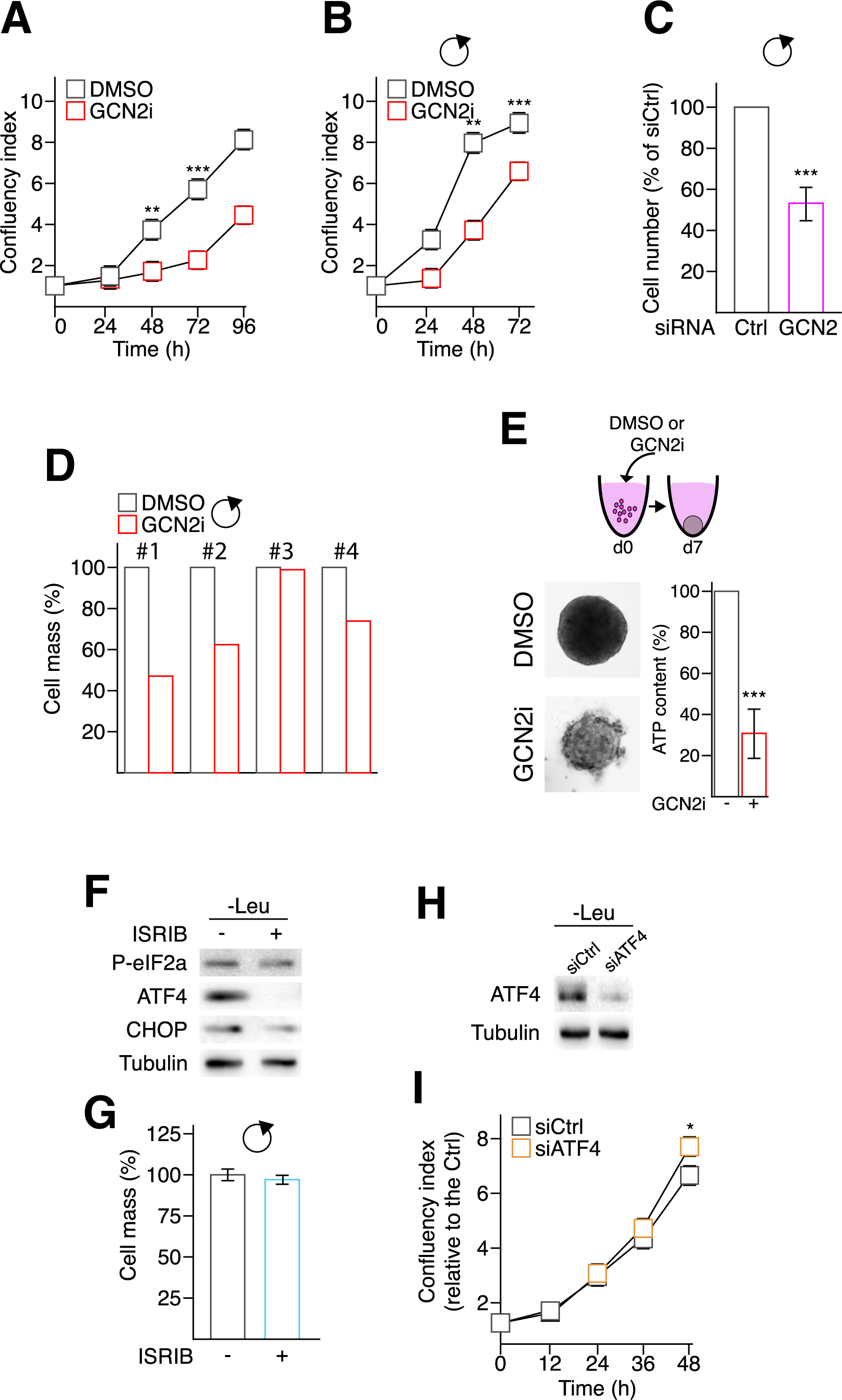
GCN2 is necessary for COAD cancer cells proliferation independently of the ISR pathway. A. Cell proliferation assessed by the confluency index of HCT116 cells treated or not with GCN2i for 96 h. Data are expressed as mean +/-s.e.m of independent experiments (n = 3). Unpaired two-tailed t-test with p-value (** p < 0.01, *** p < 0.001). B. Cell mass was assessed every day over the indicated period of time in HCT116 cells treated or not with GCN2i by SRB assay. A daily refreshment of the medium (indicated by the black rounded arrow) was performed to ensure a full supply of nutrients over the time course of the experiment. Data are expressed as mean +/-s.e.m of independent experiments (n = 3). Unpaired two-tailed t-test with p-value (** p < 0.01). C. Relative number of HCT116 cells transfected either with a siRNA control (CTRL) or against GCN2 (GCN2) following 4 days of culture. Medium were daily refreshed (black rounded arrow) to prevent nutrients exhaustion. Data are expressed as mean +/-s.e.m of 3 independent experiments. Unpaired two-tailed t-test with p-value (** p < 0.01). D. Cell mass measurement of patient-derived primary cells cultured in 2D conditions and treated with DMSO or GCN2i for 48 h. Media were refreshed daily (black rounded arrow). E. Spheroid formation assays of HCT116 treated with DMSO or GCN2i. ATP content was measured after 7 days of treatment. Treatment was initiated concomitantly to plating. Data are expressed as mean +/-s.e.m of 3 independent experiments. Unpaired two-tailed t-test with p-value (** p < 0.01). F. Western blot analysis of P-eIF2a, ATF4 and CHOP protein amounts in HCT116 starved for leucine for 8h in the presence of ISRIB (200 nM) or not. G. Measurements of cell mass in HCT116 cells treated with ISRIB (200 nM) for 48 h. Medium were daily refreshed (black rounded arrow) to ensure complete nutrients supply. Data are expressed as mean +/-s.e.m of independent experiments (n = 5). Unpaired two-tailed t-test. H. Western blot analysis of ATF4 protein levels in HCT116 cells transfected either with a siRNA control (siCtrl) or against ATF4 (siATF4) following 2 days of culture with siCTRL and siATF4 upon 8 h of amino acid deprivation (leucine starvation). I. Cell proliferation assessed by the confluency index of HCT116 cells transfected with a siCtrl and siATF4 was monitored over 48 h. Data are expressed as mean +/-s.e.m of independent experiments (n = 3). Unpaired two-tailed t-test with p-value (* p < 0.05).

To decipher the underlying mechanism by which GCN2 activity promotes cell proliferation, RNA sequencing was performed on HCT116 cells grown in 2D with daily media refreshment, silenced or not for GCN2 (Figure 5A). Gene set enrichment analysis revealed dysregulation of functions associated with protein homeostasis, particularly autophagy, and cell cycle checkpoint. Consistently, metabolomics experiments showed a perturbation in the intracellular content of several amino acids in cells silenced for GCN2 (Figure 5B) and a significant decrease in protein synthesis upon GCN2i (Figure 5C). Pharmacological inhibition of GCN2 increased the amount of the canonical autophagy markers LC3-I/II and P62 (Figure 5D). Based on the use of a molecular reporter, we confirmed the activation of an autophagic flux when the activity of GCN2 was blocked in HCT116 and HT-29 (Figure 5E, figure supplement 7A). Although treatment with a classical inhibitor of autophagy, namely chloroquine, did not affect significantly the cell mass (Figure 5F, figure supplement 7B), when both treatments were applied, chloroquine aggravated the reduction of cell mass caused by GCN2i, functionally demonstrating the protective role of the autophagic process.

**Fig. 5.**
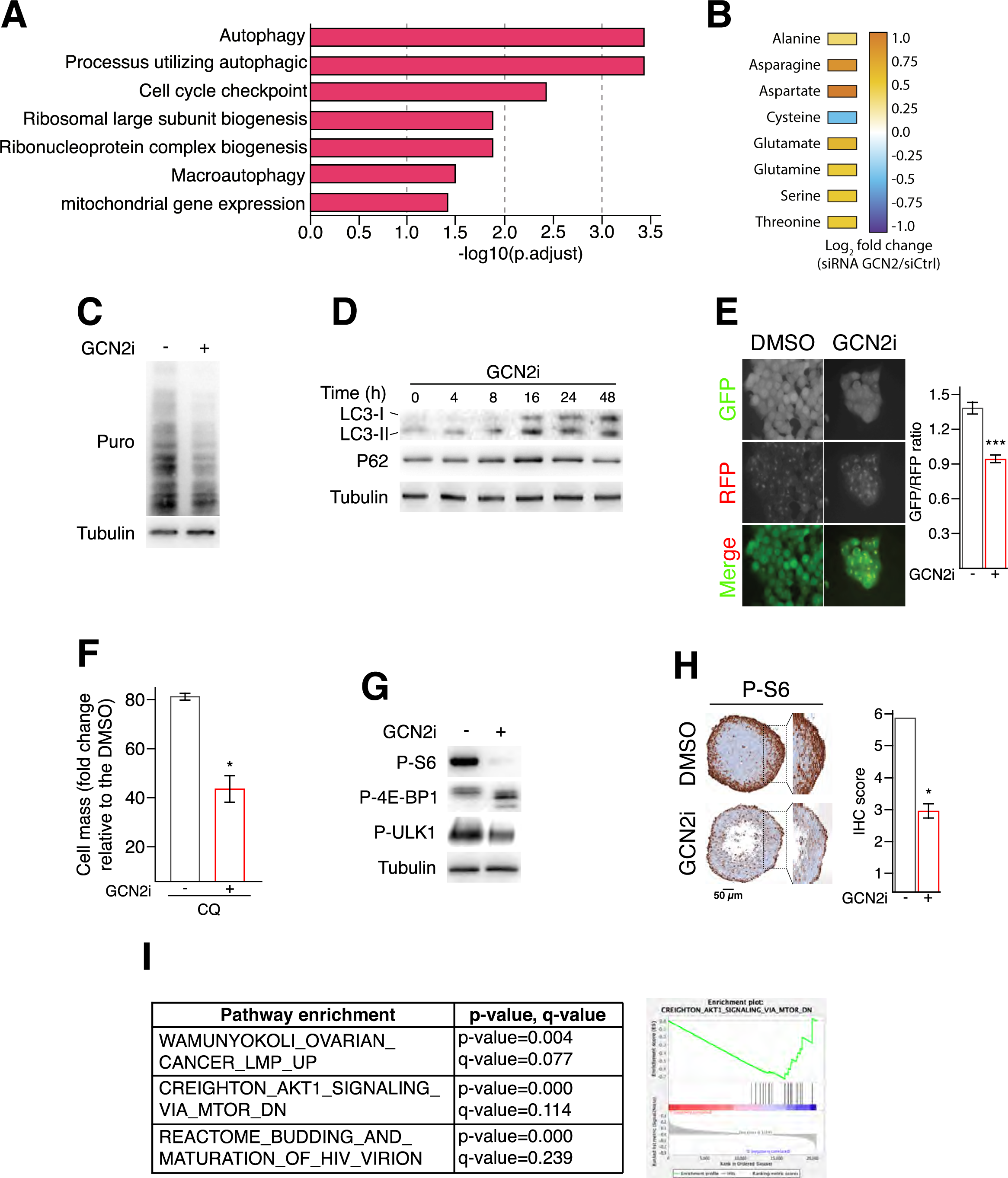
GCN2 sustains the mTORC1 activity in proliferative colon cancer cells. A. Gene set enrichment analysis in HCT116 cells silenced for GCN2 compared to the control. Top enrichment GO terms in siGCN2 transfected cells versus siCtrl are shown and significance is expressed as -Log10 adjusted p-value. B. Amino acid profiling of HCT116 cells transfected with a siRNA control or against GCN2. C. Assessment of protein synthesis rate in HCT116 cells treated for 24 h with GCN2i. D. Western blot analysis of autophagic markers, LC3-I, LC3-II and p62, in HCT116 cells treated with GCN2i for the indicated period of time. E. Autophagic flux analysis using HCT116 cells stably expressing the GFP-LC3-RFP-LC3ΔG construct and treated with GCN2i for 48 h. Cleavage of GFP-LC3 by autophagy released the RFP-LC3 as an internal control, thus reduction of the GFP/RFP ratio illustrates induction of the autophagy flux. Data are expressed as the mean of quantification +/-s.e.m. of independent experiments (n = 3). Unpaired two-tailed t test with p-value (* p < 0.05) F. Cell mass was measured in GCN2i-treated HCT116 cells after 48 h in combination with chloroquine. Data are expressed relative to the vehicle as mean +/-s.e.m of independent experiments (n = 7). Unpaired two-tailed t-test with p-value (*** p < 0.001). G. Western blot analysis of mTORC1 pathway markers (P-S6, P-4EBP1 and P-ULK1) amounts in HCT116 cells following 24 h of GCN2i treatment. H. Immunohistochemical staining against P-S6 in HCT116 spheroids treated for 3 days with DMSO or GCN2i. Quantification of the IHC staining is provided on the right panel. Data are expressed as mean +/-s.e.m of independent experiments (n = 3). Unpaired two-tailed t-test with p-value (* p < 0.05). I. Gene set enrichment analysis of COAD patients with low GCN2 expression (25%) compared to the rest of the COAD cohort (TCGA data).

We then investigated the molecular events underlying proteostasis defects when GCN2 is blocked. In complete media, mTORC1 maintains the autophagic flux at a low level and support protein translation. Additionally, previous studies report several molecular crosstalk linking mTORC1 repression to GCN2 induction or vice versa upon amino acid deprivation or rapamycin analog treatment (Averous et al., 2016; Wengrod et al., n.d.; Ye et al., 2015). Nonetheless, no evidence was reported regarding a GCN2-dependent upregulation of mTORC1 activity in proliferative state. We thus hypothesized that GCN2 inhibition could repress the mTORC1 pathway that, in turn, limits protein synthesis and induces autophagy. Activation degree of mTORC1 pathway was assessed through the amount of the phosphorylated forms of the S6 ribosomal protein, 4E-BP1 and ULK1 in COAD cells treated with GCN2i for 24 hours (Figure 5G, figure supplement 7C). The compound provoked a diminution of all tested canonical markers of mTORC1 activity, including phosphorylated ULK1 that is consistent with the autophagy activation (Kim et al., 2011). Inhibition of S6 phosphorylation by the GCN2i was also observed at the periphery of colon cancer spheroid indicating that mTORC1 pathway is also repressed in proliferating cells of 3D models and confirmed in GCN2 silenced cells (Figure 5H, figure supplement 7D). The relevance of this was then evaluated in patients using the COAD TCGA database. Our results show that this connection is independent of the ISR but is highlighted by the lack of GCN2 expression. Thus, instead of discriminating tumors according an irrelevant stress-related gene signature, we compared the low-GCN2 expressing patients to the rest of the COAD cohort in the TCGA. Gene set enrichment analysis confirmed that a low or null expression of GCN2 was significantly associated with a downregulation of the AKT-mTORC1 pathway in COAD tumors (Figure 5I).

### IV. Inhibition of GCN2 impairs MetRS translocation and 47S rRNA production

Then we sought to further define the mechanism by which lack of GCN2 activity impairs the mTORC1 signaling. RNA sequencing data indicated that a four-day GCN2 silencing triggered a dysregulation of RiBi as highlighted in figure 5A. Since RNA pol I inhibition is sufficient to repress mTORC1 and triggers subsequent autophagy, we explored whether repression of mTORC1 upon GCN2i might result from a prior defect in RiBi. This hypothesis was especially supported by the GCN2i-driven cell cycle arrest in G2/M phase that is consistent with pharmacological RNA pol I inhibition and the gene signature found in COAD patients associated to dysregulated ribosome biogenesis (Figures 1A and 6A) (Hald et al., 2019; Li et al., 2016; Ma and Pederson, 2013). Enlarged nucleoli associated with dispersion of fibrillarin were observed from 4 hours of treatment, confirming that GCN2 also controls nucleolus homeostasis in proliferative conditions (Figure 6B). Northern blot analysis revealed that GCN2 inhibition impaired 47S pre-rRNA generation (Figure 6C). However, no change in phosphorylation of S6 was observed at the same early time points, indicating that mTORC1 repression by GCN2i is rather a long-term consequence of RiBi defects upon treatment (figure supplement 8A). Then we investigated the molecular link between GCN2 and RNA pol I activity. As ISRIB treatment or ATF4 silencing did not mimic the proliferation arrest caused by the loss of GCN2 activity, a contribution of the ISR was thus excluded. However, we explored a role for the methionyl tRNA synthetase (MetRS), a second substrate of GCN2 (Kwon et al., 2011), for the two following reasons: i) MetRS has already been described to localize within the nucleolus to sustain rRNA synthesis and 47S products upon mitogenic signals although the contribution of GCN2 in this mechanism was not addressed (Ko et al., 2000) and ii) knockdown of MetRS leads to the downregulation of mTORC1 activity as well (Suh et al., 2020). Immunofluorescence assays confirmed that the colocalization of MetRS with the fibrillarin within the nucleus, in proliferative cells (figure supplement 8B). Similarly to GCN2 inhibition, MetRS silencing is sufficient to induce nucleolar stress as revealed by fibrillarin diffusion and impairs cell proliferation in nutrient-rich condition (Figure 6D, E, figure supplement 8C). We then assessed whether the cytoplasmic/nuclear shuttling of MetRS was dependent of GCN2 (Figure 6F). Inhibiting GCN2 for 2 hours of GCN2i impaired MetRS translocation. Intriguingly the GCN2iB did not affect the MetRS shuttling between the two cellular compartment. Collectively these results demonstrate that lack of GCN2 activity impairs MetRS nuclear translocation provoking a downregulation of 47S synthesis and ultimately decreased mTORC1 signaling and protein synthesis.

**Fig. 6.**
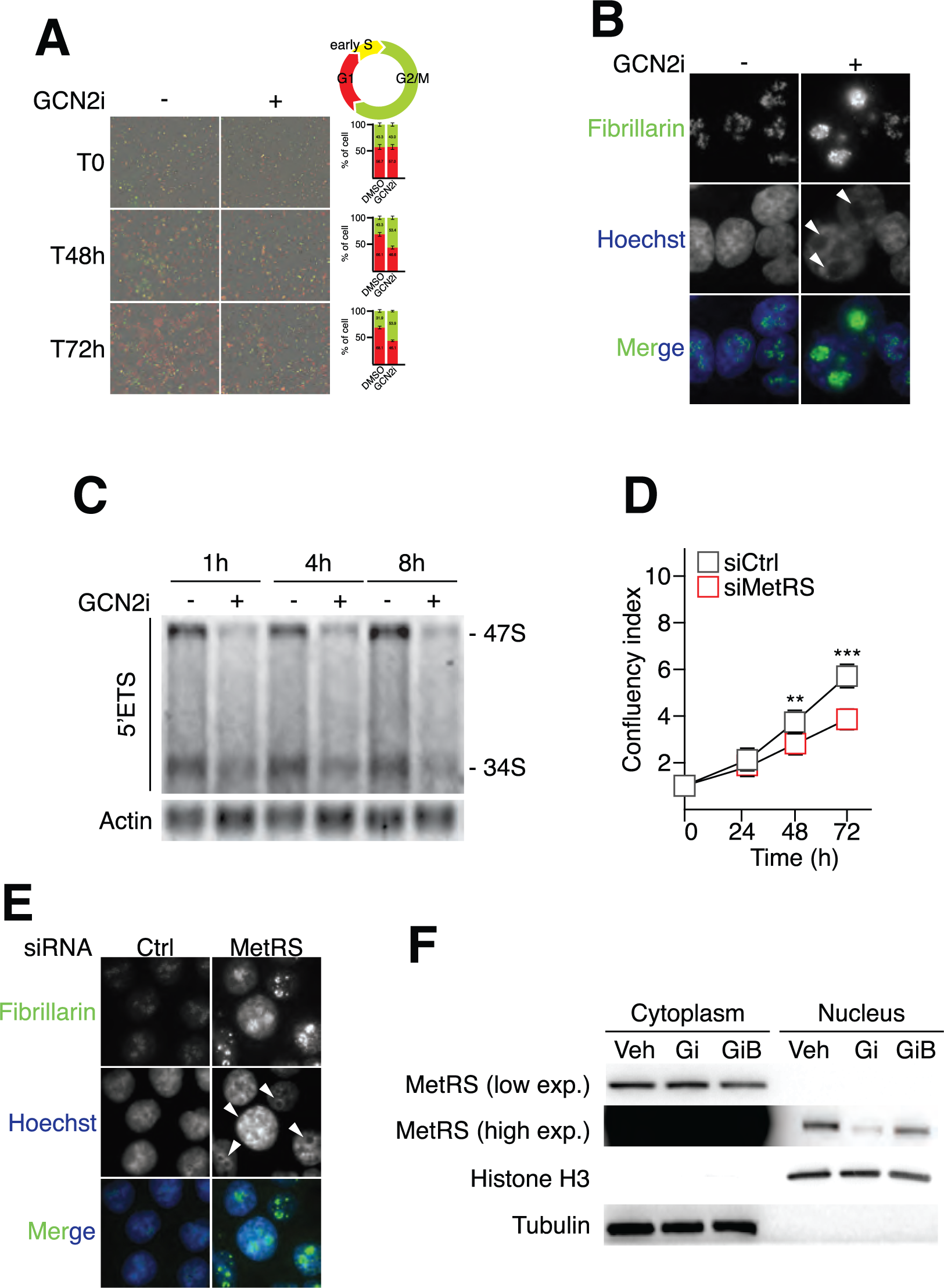
GCN2 inhibition inhibits the MetRS translocation and RNA polymerase I activity. A. Time course analysis of cell cycle progression by monitoring HCT116 FUCCI cells upon DMSO or GCN2i. Data are expressed relative to the vehicle as mean +/-s.e.m of independent experiments (n = 3). B. Fibrillarin localization assessed by immunofluorescence in HCT116 cells treated with DMSO or GCN2i for 4 hours. Nuclei were stained by Hoechst solution. C. Time course analysis of cellular rRNAs by northern blotting following GCN2i treatment. D. Cell proliferation assessed by the confluency index of HCT116 cells treated or not with GCN2i for 72 h. Data are expressed as mean +/-s.e.m of independent experiments (n = 3). Unpaired two-tailed t-test with p-value (** p < 0.01, *** p < 0.001). E. Fibrillarin localization assessed by immunofluorescence in HCT116 cells, 48 hours after transfection with a siRNA control (Ctrl) or against MetRS. Nuclei were stained by Hoechst solution. F. Western blot analysis of MetRS amounts in the cytoplasmic or nuclear fractions upon 2 hours of GCN2i and GCNiB treatments. Histone H3 and tubulin are provided as loading controls for respectively nuclear and cytoplasmic samples.

### V. Inhibition of the GCN2-MetRS axis potentiates COAD cell death upon RNA Polymerase I inhibitors in nutrient rich condition

Finally, we investigated whether impairing the GCN2/MetRS axis might confer a ribosomal vulnerability to COAD cells. We hypothesized that further inhibition of 47S pre-RNA synthesis by combining GCN2i with drugs downregulating the RNA pol I might enhance the 47S/5S disequilibrium and cause cell death. To this end, COAD cells were co-treated with GCN2i and actinomycin D, a known inhibitor of RNA pol I activity (Figure 7A) (Catez et al., 2019). After 48 hours of treatment, GCN2i or actinomycin D alone triggered a slight but significant cell death in a similar manner and their co-treatment drastically increased cell death. We then explored whether this first proof-of-principle could be relevant for translational applications. Interestingly, the mode of action of oxaliplatin, a front-line chemotherapy used in COAD treatment, also relies on the repression of RNA Pol I and induction of an irremediable ribosomal stress (Bruno et al., 2017; Sutton and DeRose, 2021). Consistently, oxaliplatin cytotoxicity is higher when cells are in the G2/M phase of the cycle, corresponding to the peak of RNA pol I activity (Klein and Grummt, 1999; Narvi et al., 2018). To test whether GCN2 inhibition may also aggravate oxaliplatin-induced nucleolar stress, immunofluorescence against fibrillarin was first performed (Figure 7B). Early formation of cap-like structures, a mark of dramatic RNA Pol I inhibition and nucleolar stress, was observed within 4 hours of administration of the drug combination. Cytotoxicity assays confirmed that GCN2i drastically increased the cell death caused by oxaliplatin in nutrient-rich conditions (Figure 7C). However, ISRIB treatment did not change the efficacy of the chemotherapeutic drug (Figure 7D). A second approach by RNA interference confirmed that downregulation of GCN2 expression specifically sensitizes COAD cells to oxaliplatin and actinomycin D (figure supplement 8D, E). Furthermore, the cytotoxicity of oxaliplatin was significantly higher in cells silenced for MetRS, (figure supplement 8F). The improvement of oxaliplatin toxicity by the inhibition of the GCN2-MetRS axis was then validated in spheroids (figure supplement 8G and supplemental movie 1). Oxaliplatin or GCN2i alone led majorly to growth impairment without clear impairment of the spheroid shape. The combination of the two compounds triggered a massive cell death at the periphery and resulted in complete unstructured spheroids. These observations were then confirmed with patient-derived colospheres in the presence or absence of the ROCK inhibitor (Figure 7F). Collectively, these results show that inhibition of the GCN2-MetRS axis drastically improves RNA pol I inhibitors-mediated COAD cell death in nutrient-rich conditions (figure supplement 8H).

**Fig. 7.**
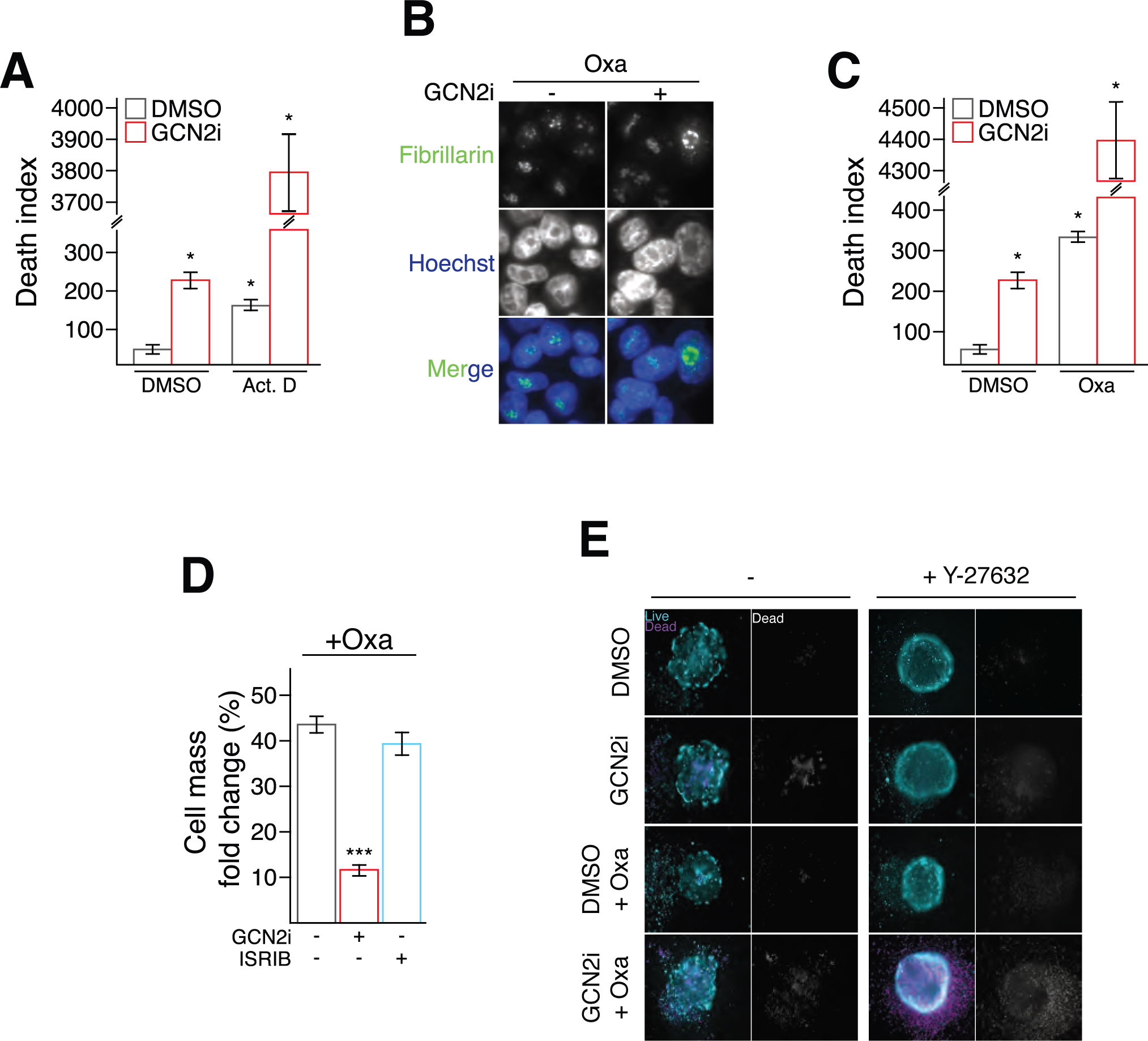
GCN2 sensitizes COAD cancer cells to inhibitors of the RNA polymerase I activity. A. Cell death index of HCT116 treated or not with GCN2i combined to Actinomycin D (Act.D) or the vehicle (DMSO) for 48 h. Data are expressed relative to the vehicle as mean +/-s.e.m of independent experiments (n = 3). One-way ANOVA with Tukey’s multiple comparisons test and p-value (* p < 0.05). B. Fibrillarin localization assessed by immunofluorescence in HCT116 cells treated with oxaliplatin (Oxa) in combination with DMSO or GCN2i for 4 hours. Nuclei were stained by Hoescht solution. C. Cell death index of HCT116 treated or not with GCN2i combined to Actinomycin D (Act.D) or the vehicle (DMSO) for 48 h. Data are expressed relative to the vehicle as mean +/-s.e.m of independent experiments (n = 3). One-way ANOVA with Tukey’s multiple comparisons test and p-value (* p < 0.05). D. Cell mass measurement of HCT116 treated for 48 h with Oxa in combination with GCN2i or ISRIB. Data are expressed relative to the vehicle as mean +/-s.e.m of independent experiments (n ≥ 3). One-way ANOVA with Tukey’s multiple comparisons test and p-value (*** p < 0.001). E. Live-and-dead assay in patient-derived colospheres, cultivated or not with ROCK inhibitor Y-27632, after 5 days of treatment with Oxa and/or GCN2i. To facilitate the observation of cell death induction, staining of non-viable cells is also represented in shades of grey.

## Discussion

Cellular plasticity relies on the integration of intrinsic and extrinsic cues at the molecular level, notably contributing to the fine tuning of the translational program. In this context, the flexibility of RiBi represents a first and critical level of cell adaptation. In this study, we identify GCN2 as a novel and determinant regulator of RiBi and nucleolar homeostasis. Intriguingly, this kinase supports tumor-cell plasticity by repressing the 47S pre-rRNA transcription upon starvation, preventing the engagement of an irreversible nucleolar stress in dedifferentiated cells. Conversely, GCN2 sustains 47S pre-rRNA transcription in nutrient-rich conditions, and the combination of inhibitors of GCN2 and RNA Pol I allows a shift from cell cycle arrest towards apoptosis.

In harsh conditions, glutamine scarcity is a major driver of tumor cell dedifferentiation leading to the acquisition of a stem cell-like phenotype associated with chemoresistance (Pan et al., 2016). The underlying molecular program is evidently complex including at least an epigenetic and translational reprogramming supported by the concomitant phosphorylation of eIF2a and repression of mTORC1 (Jewer et al., 2020). Our results show that GCN2 also supports RiBi reprogramming. The adaptive cellular response to starvation implicates limitation of both RNA Pol I activity and rRNA processing (Pan et al., 2022). Our data supports a model in which GCN2 preserves the repression of 47S expression in amino acid-depleted conditions likely to sustain the translational dedifferentiation program. These observations are consistent with the previous work showing that, upon metabolic stress, restoration of 47S pre-RNA expression through the silencing of nucleomethylin (NML) triggers cell death, mimicking our results when GCN2 activity is blocked upon starvation (Murayama et al., 2008). Whether GCN2 might influence the NML function in stressed cells has so far not been reported. Furthermore, the ATF4-mediated augmentation of Sestrin-2 in starved cells might represent the molecular link between GCN2 and 47S synthesis inhibition (Ye et al., 2015). Indeed, elevation of Sestrin-2 maintains the repression of mTORC1 upon scarcity. Furthermore, abrogation of the GCN2 activity and absence of Sestrin-2 induction triggers the mTORC1 activity. Recovery of mTORC1 activity might explain the rescue of 47S transcription in starved-cells treated with GCN2i. Inhibition of GCN2 is sufficient to further alter RiBi and trigger cell death through the TP53-proapoptotic program. Although several studies support the use of GCN2 inhibitors in the clinic to eliminate adapted cells, the mechanisms related to cell death caused by GCN2 inhibition in starved cells remain unclear. Most often attributed to an impairment of the ISR and its related translational/metabolic adaptive response, our results demonstrate that the subsequent apoptotic program likely results from the disruption of the nucleolar homeostasis. Cell death caused by the lack of GCN2 activity is accompanied by a decreased expression of c-Myc in a nutrient-scarce microenvironment. These results corroborate previous studies showing that cells overexpressing c-Myc are more sensitive to GCN2 inhibition (Schmidt et al., 2019; Tameire et al., 2019). Thus, one can postulate that increased c-Myc expression stimulates the glutaminolysis pathway and aggravates microenvironmental exhaustion of Gln. As a consequence of the c-Myc-driven nutritional stress, GCN2 activation and subsequent RiBi downregulation preserve the nucleolus homeostasis and optimize energy efficiency (Gao et al., 2009; Shore and Albert, 2022; Wise et al., 2008). Nonetheless, cell death caused by GCN2 inhibition is also effective in models expressing a wild type form of APC (e.g. HCT116), classically associated with a lower expression of c-Myc in COAD. These results suggest that, beyond the mutational status of APC previously proposed, the subset of tumors expressing a wild-type form of TP53 should be prioritized, considering that high levels of c-Myc might provide an optimal response to GCN2 inhibitors. Moreover, although GCN2 activity is required for repressing 47S expression upon low glutamine, whether the ISR axis plays a critical role in this mechanism requires further investigations. The Dr. Lyons’ group recently showed that chemical stress (NaAsO_2_), activating the HRI-p-eIF2a axis, limits the first step of rRNA processing. However, the authors demonstrated that this effect is independent of both HRI and eIF2a suggesting that, upon stress, an ISR-independent pathway controls the RiBi pathway (Szaflarski et al., 2022). Beside HRI, GCN2 is also activated in response to arsenate (Taniuchi et al., 2016). Thus we propose that, instead of HRI, GCN2 might be responsible of the modulation of the RiBi and determine the cell outcome upon NaAsO_2_ (Taniuchi et al., 2016). Collectively, these results support the hypothesis that unknown substrates of GCN2 may be at the interface between external stresses and RiBi.

In addition to the canonical function of GCN2 in stressed cells, we found that this kinase also functions in nutrient-rich conditions. Our robust experimental set-up, distinguishing starved and non-starved conditions, revealed that inhibition of GCN2 represses proliferation and the mTORC1 pathway. This effect appeared to be cell type-dependent since proliferative DLD-1 or primary cells isolated from one COAD patient were insensitive to the loss of GCN2 activity. It is likely that mTORC1 repression is a consequence of RiBi impairment following GCN2 inhibition. Indeed, alteration of 47S synthesis is an early event in refreshed medium settings and is not associated, at these early time points with impaired phosphorylation of S6. These observations reinforce the notion that GCN2 also controls RiBi in the absence of nutritional stress. The mechanism underlying this novel cell-autonomous function for GCN2 implicates its capacity to sustain MetRS translocation within the nucleus to ensure the stimulation of RNA Pol I and cell proliferation. How MetRS controls the 47S transcription and largely RiBi remains unknown. Yet, in line with our data, high expression of MetRS is of poor prognosis in several types of human tumors, including COAD, and consistently associated to enhanced mTORC1 activity and cancer cell proliferation confirming its role in promoting tumor growth (Jin et al., 2020; Kim et al., 2017; Kushner et al., 1976).

Finally, this work shows that GCN2 represents a novel regulator of the RiBi in deprived and non-deprived cells that can be therapeutically exploited in the clinic through the use of compounds targeting the kinase. Overall, the threshold of repression of the RNA Pol I activity is a key determinant in the induction of cell death. In this context, gastrointestinal cancers may constitute interesting types of cancer models since they are classically treated using FOLFOX or FOLFIRI, two chemotherapeutic regimens containing drugs targeting the ribosomal function. Indeed, oxaliplatin was reported to kill cells through the induction of a ribosomal stress and 5-fluorouracil was described to impair 47S processing and recently to be integrated in the ribosome and promoting profound translational reprogramming (Bruno et al., 2017; Burger et al., 2010; Ge et al., 2017; Sutton and DeRose, 2021; Therizols et al., 2022). Nonetheless, these compounds are mainly effective on proliferative cells since metabolic stress limits their efficacy (Xu et al., 2019). Moreover, glutamine starvation drives resistances to RNA Pol I inhibitors. Our results provide a solid basis for promoting the transfer of GCN2 inhibitors to the clinic as a promising approach and argue in favor of developing compounds inhibiting concomitantly the ISR and MetRS branches of the pathway and satisfying all the physicochemical requirements for in vivo applications. Such a strategy would allow: i) elimination of chemoresistance of TP53-WT quiescent tumor cells, and ii) sensitization of proliferative cells towards oxaliplatin or other RNA pol I inhibitors. The benefits of in such inhibitors would be of a greater translational value than ISR-centered compounds currently under development.

## Materials and methods

### Cell culture

The following cell lines were obtained from ATCC: HCT116, HT29, DLD1 and LoVo. HCT116 TP53 WT and HCT116 TP53 KO were generated previously by Dr B. Vogelstein, Ludwig Center at Johns Hopkins, Baltimore (Bunz *et al*, 1998) and were kindly provided by Drs P. Mehlen and A. Paradisi at CRCL, Lyon. HCT116 FUCCI were obtained by transducing HCT116 with the Cdt1-mKO and Geminin-mAG constructs and FACS cell sorting. All cell lines were routinely cultured in McCoy medium (ref 16600082, GIBCO, Thermo Fisher Scientific, Whaltam, Massachussets, USA), except DLD1 cultured in RPMI medium (ref 21875034, GIBCO, Thermo Fisher Scientific, Whaltam, Massachussets, USA). Cell lines were tested for absence of mycoplasma after amplification. All media were supplemented with 10% fetal bovine serum (FBS) and 1% v/v penicillin/streptomycin (ref 15140122, GIBCO, Thermo Fisher Scientific, Whaltam, Massachussets, USA). Cells were maintained at 37°C in a 5% CO_2_ incubator. For 2D experiments, treatment was initiated the day after plating with daily refreshment when indicated by a circular arrow. Glasstic slides (ref 87144, Kova International, Garden Grove, California, USA) were used for cell counting.

For patient-derived tissue culture, after tumor dissociation using a mix of DNAse and collagenase, individualized tumor cells were embedded in Matrigel matrix (ref 354234, Corning, New York, USA) and cultivated as tumoroids using the Stemness system device and IntestiCult Organoid Growth Medium (ref 06010, Stemcell, Saint Égrève, France). The collection and use of all patient tissue specimens was carried out according to French laws and regulations in the frame of the ColonIM protocol (NCT03841799).

### Nutritional manipulation and reagents

Leucine starvation experiments were performed using a DMEM medium devoid of leucine, lysine and arginine (ref 88425, GIBCO, Thermo Fisher Scientific, Whaltam, Massachussets, USA), in which lysine (final concentration 0.8 mM) (ref L8662, Sigma Aldrich, Saint Louis, Missouri, USA), arginine (final concentration 0.4 mM) (ref A6969, Sigma Aldrich, Saint Louis, Missouri, USA), non-essential amino acids (ref 11140050, GIBCO, Thermo Fisher Scientific, Whaltam, Massachussets, USA), 1% penicillin/streptomycin (ref 15140122, GIBCO, Thermo Fisher Scientific, Whaltam, Massachussets, USA) and 10% dialyzed serum were subsequently added. The control medium for these experiments was supplemented with leucine (final concentration 0.45 mM) (ref L8912, Sigma Aldrich, Saint Louis, Missouri, USA)

To perform experiments in condition of low asparagine or low glutamine medium, we used Dulbecco’s Modified Eagle Medium devoid of glucose and sodium pyruvate (ref A1443001, GIBCO, Thermo Fisher Scientific, Whaltam, Massachussets, USA), supplemented with 1% penicillin/streptomycin (ref 15140122, GIBCO, Thermo Fisher Scientific, Whaltam, Massachussets, USA), 1% sodium pyruvate (ref 11360070, GIBCO, Thermo Fisher Scientific, Whaltam, Massachussets, USA), and 10% dialyzed FBS. The concentrations mentioned in Tajan et al. (Tajan *et al*, 2018) were used to supplement control medium with glucose (final concentration 16 mM) (ref A2494001, GIBCO, Thermo Fisher Scientific, Whaltam, Massachussets, USA) and the subsequent amino acids purchased as powders from Sigma Aldrich (Saint Louis, Missouri, USA) : L-proline (0.15 mM, ref P5607), L-alanine (0.15 mM, ref 05129), L-aspartic acid (0.15 mM, ref A8949), L-glutamic acid (0.03 mM, ref W328502), L-asparagine (0.34 mM, ref A4159) and L-glutamine (2 mM, ref G3126). For nutrient scarce medium, L-asparagine and L-glutamine were added respectively at the concentrations of 0.02 mM and 0.1 mM based on the work of Pan et al (Pan *et al*, 2016).

GCN2 inhibitor (GCN2i) used this study was previously described as TAP20 in (Dorsch *et al*, 2014; Bröer *et al*, 2019) and generously provided by Merck KGaA. The compound was resuspended in DMSO and, if not mentioned otherwise, used at a final concentration of 5 µM. The GCN2iB was purchased from MedChemExpress (Monmouth Junction, NJ) and also used at final concentration of 5 µM. Oxaliplatin was provided by the Centre Léon Bérard pharmacy, diluted in H_2_O and used at a final concentration of 5 µM. ISRIB (ref SML0843, Sigma Aldrich, Saint Louis, Missouri, USA) dissolved in DMSO was used at a final concentration of 200 nM. Chloroquine (ref C6628, Sigma Aldrich, Saint Louis, Missouri, USA) was prepared in PBS pH5 and used at a final concentration of 20 µM. Rapamycin (ref T1537, Euromedex, Souffelweyersheim, France) dissolved in DMSO was used at a final concentration of 10 nM. Actinomycin D (ref A1410, Sigma Aldrich, Saint Louis, Missouri, USA) was diluted in DMSO and treatments were performed at a final concentration of 1 nM.

Regarding RNA interference experiments, HCT116 cells were seeded onto 6-well plates (2.5 × 10^5^ cells per well) and concomitantly transfected with non-targeting siRNA (sc-37007) or human GCN2 siRNA (sc-45644) or human ATF4 siRNA (sc-35112) or human MetRS siRNA (sc-75775) (Santa Cruz Biotechnology, Dallas, Texas, USA). All siRNA transfections were performed using 50 nM siRNA and HiPerFect reagent (ref 301704, Qiagen, Hilden, Germany), according to the manufacturer’s traditional transfection protocol. Day after transfection, media were changed and treatments were performed 48 h later.

### Cell cycle progression, Proliferation and Cytotoxicity assay in two-dimensional models

For cell cycle progression, HCT116 FUCCI cells were plated onto 6-well plate (2.50 x 10^5^ cells per well). The green and red fluorescence of the FUCCI reporter, indicative of the cell cycle phase, was monitored and quantified over time using the Cellcyte X system (Cytena, Freiburg im Breisgau, Germany). Then, the percentages of green and red fluorescence were estimated in each studied condition to respectively evaluate the proportion of cells in G1 or G2/M phases of the cell cycle.

For proliferation and cell death assessment in 2D culture, colon cancer cells (1x10^5^ cells/well) were seeded onto a 24-well plate and subjected 24h later to the indicated treatment. Propidium iodide was added in the media at 2.5 µg/mL final for apoptosis assay. Inhibitors of apoptosis Q-VD-OPh (QVD 20 µM, ref 1170-3) and Z-VAD-FMK (ZVAD, 20 µM, ref A12373-5), and inhibitors of necroptosis necrostatin-1 (Nec1, 10 µM, ref A4213) and RIPK3 inhibitor (R3i, 5 µM, ref 2673-5) were all purchased from Clinisciences (Nanterre, France). Then, cell confluency and cell death were monitored over the indicated period of time with data collection every hour using the Cellcyte X system (Cytena, Freiburg im Breisgau, Germany). Death index was calculated using the formula “(PI Foci Number/Confluency) x100”.

Cell mass was assessed by using the sulforhodamine B (SRB) assay. Cells were seeded onto 6-well plates (2.50 × 10^5^ cells per well) and treated the day after plating for 24 to 96 h at 37°C, 5% CO_2_ depending on the experiment. After removing the medium, wells were washed with PBS. Then, 1 mL of 10% trichloroacetic acid was added and after 1 h of incubation at 4°C, the plates were flicked and washed three times with tap water before being dried at room temperature for 1 h. The wells were then stained with 1.5 mL of SRB solution 0.057% in 1% acetic acid (ref 341738, Sigma Aldrich, Saint Louis, Missouri, USA) for 30 min at room temperature under gentle agitation. Plates were flicked and washed three times with 1% acetic acid and dried at 37°C for 1 h. Finally, 1.5 mL of 10 mM Tris base was added and shaken vigorously for 15 min. The absorbance was measured using Tecan Infinite M200 Pro (Tecan, Männedorf, Switzerland) at a wavelength of 510 nm.

### Autophagic flux monitoring

To monitor autophagic flux, 293GP cells were transfected with a pMRX-IP-GFP-LC3-RFP-LC3ΔG construct described in Kaizuka et al (Kaizuka *et al*, 2016). Then, viral supernatant was collected to infect HCT116 and HT29 cells and generate stable cell lines carrying pMRX-IP-GFP-LC3-RFP-LC3ΔG plasmid, subsequently renamed HCT116 pLC3 and HT29 pLC3. Cells were then grown on glass coverslips with DMSO or GCN2i for 48h. After that, cells were fixed in 4% of paraformaldehyde in phosphate-buffered saline (PBS). Coverslips were mounted on microscope slides using the Fluoromount G mounting medium (ref 17984-25, EMS Diasum, Hatfield, Pennsylvania, USA). Nikon ECLIPSE NI-E microscope with a 100X immersion objective (Nikon, Tokyo, Japon) was used to acquire images. Final images were analyzed and cropped with the Fiji/ImageJ software (National Institutes of Health, Bethesda, Maryland, USA).

### Spheroid and tumoroid assays

For spheroid formation assays, drops of 10µL containing 3 x 10^3^ cells were deposed on the lid of plastic dishes. The lid was inverted and placed over the dish containing 5mL of sterile PBS in the bottom. Treatment with GCN2i is concomitant to plating. Hanging drops were then maintained at 37°C in a 5% CO_2_ incubator for 7 days.

For Live-and-Dead cell assays, 3 x 10^3^ cells from patient primary cells or indicated cell lines were plated per well in 96-wells ultra-low attachment plates (ref 650970, Greiner Bio-One, Les Ulis) 3 days before treatment. After 72 hours of treatment, viable and dead cells were stained using the Live/Dead cell imaging kit provided by Thermo Fisher (ref R37601, Thermo Fisher Scientific, Whaltam, Massachussets, USA) according to manufacturer’s instructions. Fluorescence microscope was used to capture fluorescence signaling and then, pictures were colored and cropped with the Fiji/ImageJ software (National Institutes of Health, Bethesda, Maryland, USA).

### Amino acid profiling

Before analysis, a mix of 24 internal standards was added to cell extracts. After homogenization and centrifugation, the supernatant was evaporated under nitrogen and the dry residue was resuspended in 200 µL mobile phase. Separation was performed by liquid chromatography on a Ultimate 3000 system (ThermoFisher Scientific™, Bremen, Germany), using a Synergi HydroRP column (250 x 2; 4 µm, Phenomenex) and 0.1 % of formic acid in water and methanol as mobile phases. The detection was performed with a Q-Exactive Plus Orbitrap mass spectrometer (ThermoFisher Scientific™, Bremen, Germany) equipped with a heated electrospray ionization source which operated in positive and negative modes. Retention time, exact mass and isotopic pattern of the molecular ion, exact mass of the fragments and comparison of the fragmentation spectrum with a homemade database were used for identification of metabolites. The ratio (peak area of metabolites / peak area of internal standard) was used for quantification.

### RNA extraction and RT-qPCR

Total cellular RNA was extracted after the indicated period of treatment using TRIzol Reagent (ref 15596026, Invitrogen, Carlsbad, California, USA) according to the manufacturer’s protocol. For cDNA synthesis, 0.5 µg of RNA were reverse transcribed using Superscript II reverse transcriptase (ref 18064014, Invitrogen, Carlsbad, California, USA) with random primers (S0142, Thermo Fisher Scientific, Whaltam, Massachussets, USA), according to the manufacturer’s instructions. cDNA was then amplified by qPCR using specific primers listed in supplementary Table 1 and the SYBR Green Master Mix (ref 1725274, Bio-Rad, Hercules, California, USA). qPCR was performed using the CFX connect real-time PCR system (Bio-Rad, Hercules, California, USA). Relative quantification was assessed using a standard curve-based method. Expression of target genes was normalized against RPS11 mRNA levels used as an internal control. qPCR experiments were repeated at least three times in duplicate.

### Western blot analysis and SUnSET assays

To perform Western blot analysis, cells were seeded onto 100mm-diameter plates (1.50 × 10^6^ cells per plate) and treated for the incubation period of corresponding figures. Whole cell extracts were prepared from cultured cells lysed at 4°C in RIPA protein buffer containing protease and phosphatase inhibitors (ref 11697498001, Roche, Bâle, Switzerland), and obtained by centrifugation at 13,000 × g for 20 min at 4°C. Subcellular fractionation was performed as follows: cells were scratched on ice and collected in cold PBS before centrifugation at 3,000 x g for 5 min à 4°C. The pellet was resuspended and lysed on ice in a hypotonic buffer (10 mM Hepes pH 7.5, 10 mM KCl, 0.1 mM EDTA, 1 mM DTT, 0.5% NP-40 and protease/phosphatases inhibitors) for 15 min. Centrifugation was performed at 12,000 x g for 5 min at 4°C and the supernatant containing the cytoplasmic fraction was collected and an equal volume of Laemmli protein buffer 2X containing protease and phosphatase inhibitors was added. The pellet, representing the nuclear fraction, was dissolved in the hypotonic buffer and centrifugated again at 12,000 x g for 5 min at 4°C. The supernatant was removed and the pellet containing the nuclear fraction was resuspended in Laemmli protein buffer 1X supplemented with protease and phosphatase inhibitors. The nuclear fraction was obtained by centrifugation at 1500 × g for 30 minutes at 4°C. Tubulin and Histone H3 were used as loading control for purity of cytoplasmic and nuclear fractions, respectively. Protein concentrations of the cellular extracts were determined using the DC Protein Assay (ref 5000112, Bio-Rad, Hercules, California, USA). Equal amounts of proteins (20 μg) were separated by SDS-PAGE and then transferred onto nitrocellulose membranes (ref 1704271, Bio-Rad, Hercules, California, USA). Membranes were incubated in blocking buffer, 5% milk or Bovine Serum Albumin (BSA) in Tris-Buffered Saline/Tween 20 (TBST), for 1 h at room temperature, then incubated overnight at 4°C with the appropriate primary antibodies diluted in TBST containing 5% milk or BSA. Membranes were washed three times with TBST, incubated for 1 h at room temperature with the appropriate secondary antibodies, diluted in TBST containing 5% milk, and again washed three times with TBST. Detection by enhanced chemiluminescence was performed using the Immobilon Forte HRP Western substrate (ref WBLUF0500, Merck Millipore, Darmstadt, Germany) or SuperSignal West Femto Maximum Sensitivity Substrate (ref 34096, Thermo Scientific, Thermo Fisher Scientific, Whaltam, Massachussets, USA). Tubulin was used as a loading control. The primary antibodies used were purchased either from Santa Cruz Biotechnology (Dallas, Texas, USA): TP53 (sc-47698) and CHOP (sc-7351); from Cell Signaling Technology (Danvers, Massachussets, USA): Tubulin (2146), Histone H3 (4499), BiP (3177), P-eIF2a (3398), ATF4 (11815), LC3 (2775), P62 (5114), P-S6 (4858), P-4EBP1 (2855), P-Ulk1 (14202) and GCN2 (3302); from Merck Millipore (Darmstadt, Germany): Puromycin clone 12D10 (MABE343); from ABclonal Technology (Woburn, Massachussets, USA): MetRS (A9938); or from Calbiochem (San Diego, California, USA): MDM2 clone 4B2C1.11 (OP145). The HRP-conjugated secondary antibodies (anti-rabbit and anti-mouse antibodies, respectively ref 7074 and 7076) were supplied by Cell Signaling Technologies (Danvers, Massachussets, USA).

Rates of nascent protein synthesis were evaluated using the surface sensing of translation (SUnSET) method as previously described by Schmidt et al. *(Schmidt* et al*, 2009)*. Cells were seeded onto 6-well plates (2.50 × 10^5^ cells per well) and treated the day after plating with GCN2i for 48 h at 37°C, 5% CO_2_. 15 min before being harvested and processed to prepare whole cell extracts in RIPA buffer, cells were incubated with 5 μg/mL of puromycin (ref P9620, Sigma Aldrich, Saint Louis, Missouri, USA) directly added into the medium. The amount of puromycin incorporated into nascent peptides was then evaluated by Western blot analysis on 20 μg of proteins using anti-puromycin antibody (ref MABE343) purchased from Merck Millipore (Darmstadt, Germany).

### Immunohistochemistry and immunofluorescence staining

HCT116 cells were plated (3 x 10^3^ cells/well) and grown for 3 days in 96-wells ultra-low attachment plates (ref 650970, Greiner Bio-One, Les Ulis) before treatments. 72h after being treated, samples were formalin-fixed, embedded in paraffin and stained with antibodies purchased at Cell Signaling Technology (Danvers, Massachussets, USA): cleaved caspase 3 (9664), cleaved PARP (5625), P-S6 (4858). Immunohistological analyses were performed at the Plateforme Anatomopathologie Recherche of the Centre de recherche en cancérologie de Lyon (CRCL).

Colon Disease spectrum tissue microarray (TMA) was obtained from US Biomax (ref BC5002b, Derwood, Maryland, USA) as fresh-cut slides. The tissue microarray containing Colon Malignant and Normal Adjacent Tissue cores was stained against GCN2 (Cell Signaling Technology, 3302) by immunohistochemistry. The analysis of the IHC-stained TMA slide was performed using QuPath0.3.2 software (Bankhead *et al*, 2017). Slides were scanned and imported into the program. Positive cells detection was performed following QuPath established method and then, rigorous quality control (QC) steps were taken to remove necrosis or keratin and tissue folds. IHC staining was scored for each TMA core using the built-in scoring tool. This analysis was double-checked by an anatomopathologist.

For immunofluorescence assays, cells were grown on glass coverslips, fixed in 4% of paraformaldehyde in phosphate-buffered saline (PBS) before permeabilization with 0.5% Triton X-100 in PBS. Fibrillarin were detected using the anti-FBL mouse polyclonal antibody (ref ab4566, Abcam, Cambridge, England) diluted at 1:500. Secondary Antibody Alexa Fluor 488 Goat anti-mouse IgG (ref A-11029, Life Technologies, Carlsbad, California, USA) was used at 1:800. Nucleus were stained for 5 min with Hoechst n°33342 (ref H3570, Invitrogen, Carlsbad, California, USA) at 1:2000. Coverslips were mounted using the Fluoromount G mounting medium (ref 17984-25, EMS Diasum, Hatfield, Pennsylvania, USA). Nikon ECLIPSE NI-E microscope with a 100X immersion objective (Nikon, Tokyo, Japon) was used to acquire images. Final images were analyzed and cropped with the Fiji/ImageJ software (National Institutes of Health, Bethesda, Maryland, USA).

### Northern blot analyses

For rRNA transcription/processing analysis, HCT116 cells were cultivated with treatments as previously described for the indicated period. Then, total RNA was extracted with TRIzol according to manufacturer’s instruction as described before. Purified RNAs were resuspended in water and quantified by spectrophotometry. Briefly, 10 or 5 µg of total RNA was denaturated in formamide loading buffer and electrophoresed in denaturating agarose gel. Then, separated RNAs were transferred and crosslinked on a nylon zeta-probe membrane before hybridization with 5’ETS probes coupled with infrared-emitting fluorophores (sequences provided in supplementary table 3). Blots were acquired using Chemidoc MP Imaging System and analyzed with ImageLab software.

### RNA sequencing

For RNA sequencing, HCT116 cells were transfected with a control siRNA or human GCN2 siRNA as previously described. RNA was extracted 48h later using NucleoSpin RNA kit (ref 740955, Macherey-Nagel, Düren, Germany). Library preparation and RNA sequencing were performed at the ProfileXpert platform (UCBL, Lyon). Quality of samples was checked by Bioanalyzer 2100 (Agilent technologies, Santa Clara, California, USA) and RNA was quantified by Quantifluor RNA kit (Promega, Madison, Wisconsin, USA). First, mRNA was enriched from 1 µg of total RNA, then library preparation was realized with the NextFlex Rapid Directional mRNA-Seq kit (Bioo-Scientific, Perkin-Elmer, Walthman, Massachusetts, USA). Quality of libraries were checked by Fragment Analyzer (Agilent technologies, Santa Clara, California, USA) and quantified by qPCR. Samples were put on Flow Cell High Output. Amplification and sequencing were performed with Illumina NextSeq500: run Single Read 76 bp was performed. After demultiplexing of the data with Bcl2fastq v2.17.1.14, trimming was performed using cutadapt v1.9.1 software. Then the reads were mapped using Tophat2 v2.1.1 software with default parameters on the human genome GRCh38. The Fragments Per Kilobases of exon per Million mapped reads (FPKM) values were then computed for each gene using Cufflinks v2.1.1 software. Genes with a FPKM inferior to 0.01 were substituted by 0.01. Differential expression was estimated with a t-test with unequal variance and p-value were corrected by Benjamini and Hochberg unsing the False Discovery Rate (FDR) controlling procedure. Genes were considered as differentially expressed if the absolute value was greater than or equal to 1.5 and p-value was less than 0.05. For each comparison, transcript was considered if the coverage was greater than or equal to 1 for all samples of at least one group. VolcanoR was used to perform the enrichment analysis (Naumov *et al*, 2017).

### Patients’ survival and expression data analyses

Disease-specific survival were determined by using the Long-term Outcome and Gene Expression Profiling Database of pan-cancers (http://bioinfo.henu.edu.cn/DatabaseList.jsp) and the TCGA COAD cohort. GCN2 expression and respective statistical analyses were performed by using: the Cancertool interface (Cortazar *et al*, 2018) for Marisa et al. data (Marisa *et al*, 2013), UALCAN (Chandrashekar *et al*, 2017) for analyzing expression of GCN2 in COAD according to the stage of the disease (TCGA data), Colonomics (https://www.colonomics.org/expression-browser) for the paired samples comparison. USCS Xena Browser (Goldman *et al*, 2020) was used to perform Gene Ontology enrichment analysis on the TCGA database, comparison of genes expression and correlation between genes signatures. Enrichement of gene ontology analyses were performed by using the application Enrichr (Kuleshov *et al*, 2016).

### Statistics

All of the statistical analyses were performed using GraphPad Prism version 6.0 (GraphPad Software, San Diego, CA, USA) by using unpaired t-test or one-way ANOVA. Survival analysis was performed by Kaplan–Meier and Log-rank test. p-values less than 0.05 were considered statistically significant.

## Acknowledgements

We thank Drs. L. Le Cam and P. Fafournoux for their insightful comments sharing material and manuscript improvement, B. Manship for English editing, Y. Chaix for funding management. The authors are grateful to the members of the Anatomopathologie Recherche Lyon-Est, Imagerie Cellulaire, ProfileXpert and AniPath facilities for their collaborative work. This work was supported by the Cancéropôle CLARA (CVPPRCAN000174, CVPPRCAB000180 and CV-2021-039), Region Auvergne Rhone-Alpes (19-010898-01), Projets Fondation and Aide doctorale (R16173CC, ARCMD-Doc22021020003295) from ARC, Ligue Nationale contre le Cancer (R17167CC, R19007CC), Institut Convergence François Rabelais (17IA66ANR-PLASCAN-MEHLEN), the Marie Sklodowska-Curie fellowship (n_839398), CNRS Prematuration (NA-7-07-20), and the IPR (Innovation Pharmaceutique et Recherche) program.

