## Supplementary data for "Targeting the cell and non-cell autonomous regulation of 47S synthesis by GCN2 in colon cancer"

**Affiliations:**

Supplementary information includes:

- Figure supplement S1-S8
- Supplementary tables 1-3
- Supplementary movie 1

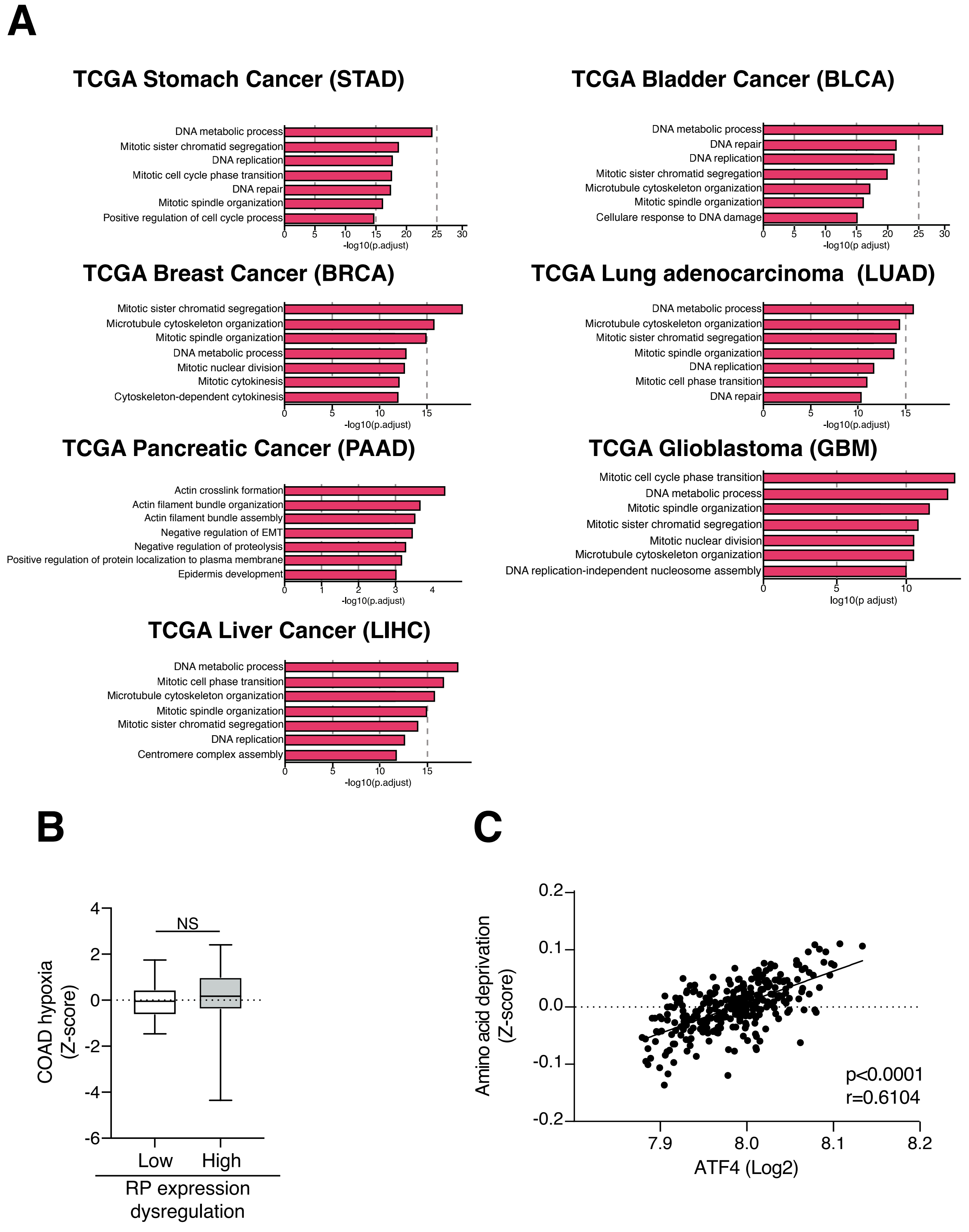

**Figure supplement 1. Enrichment assessment of gene sets and signatures of interest in various types of cancers.**

A. Gene Ontology enrichment analysis on TCGA database of different cancers.

B. Comparison of the COAD hypoxia gene signature in COAD cohorts with low and high levels of dysregulated riboproteins expression.

C. Correlation between amino acid deprivation and ATF4 gene signatures.

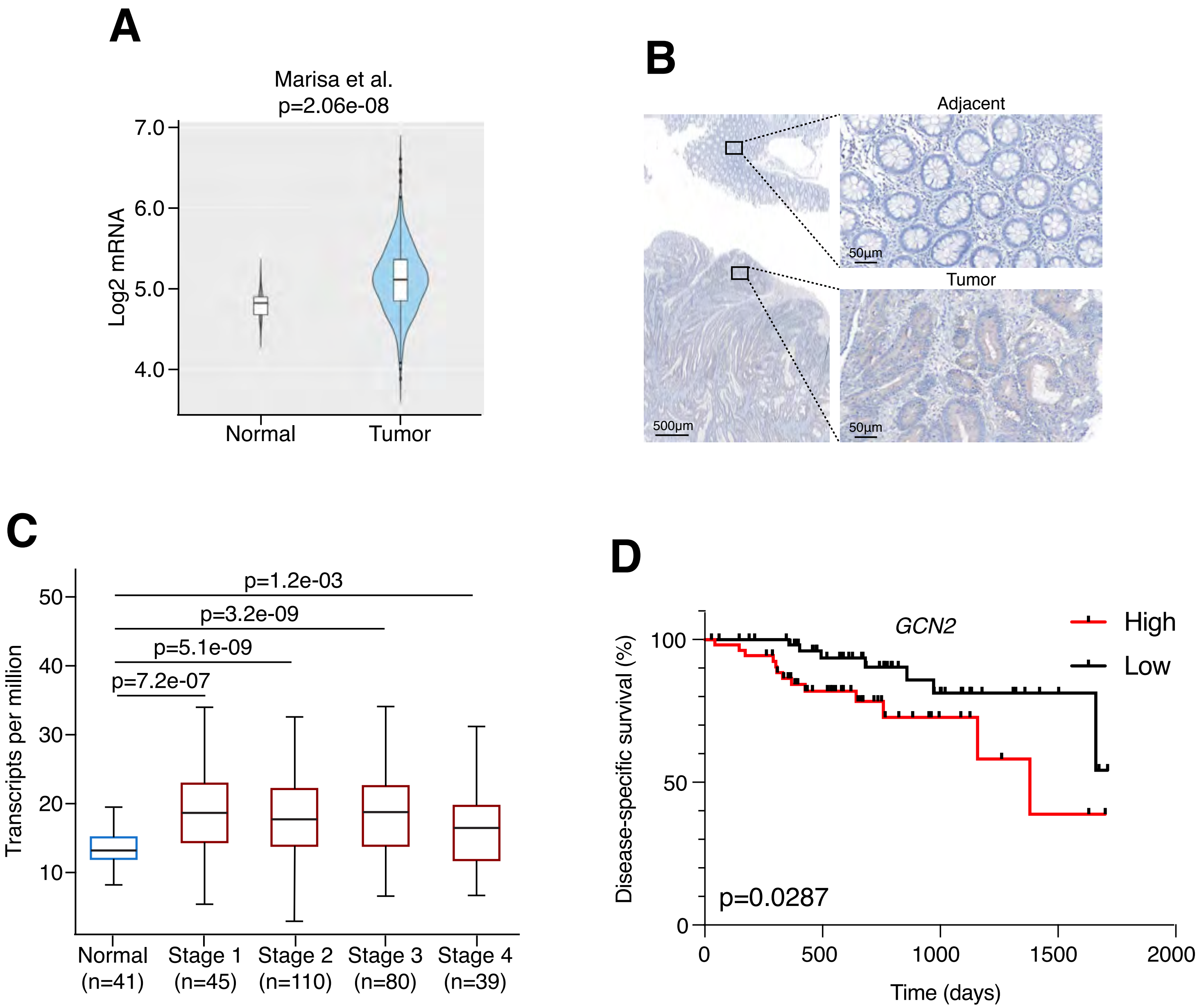

**Figure supplement 2. GCN2 accumulation and prognostic value in human colon adenocarcinoma.**

A. GCN2 mRNA levels in normal tissue compared to tumoral tissue. (Marisa et al. cohort)

B. Example of GCN2 amount accumulation in the tumor cells compared to the normal adjacent tissue.

C. Analysis of GCN2 expression accross normal tissue and the 4 stages of colon adenocarcinoma (TCGA database).

D. Disease specific survival of COAD patients with high (red) or low (black) expression of GCN2 5 years after the diagnosis (TCGA data).

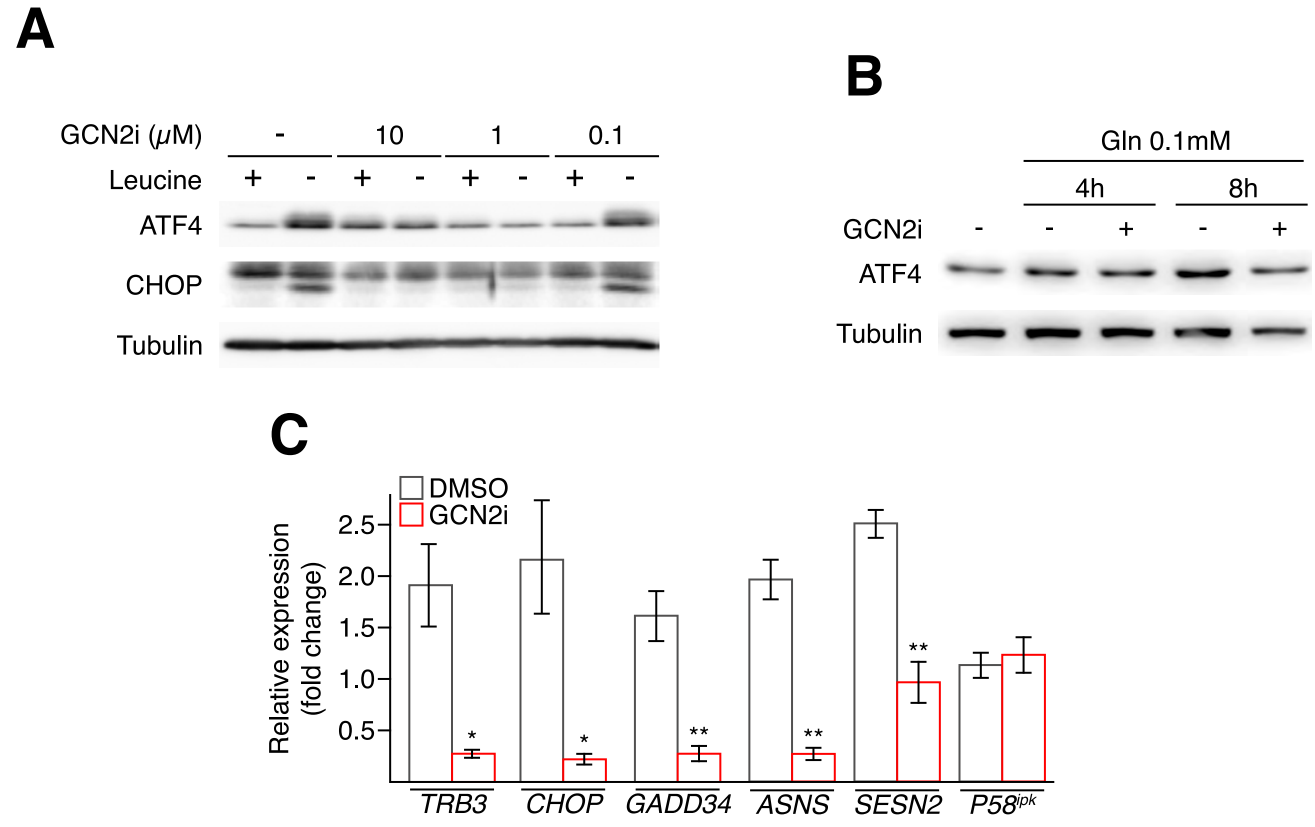

**Figure supplement 3. Validation of the ISR impairment by the GCN2i.**

A. Western blot analysis of ISR markers induction (ATF4 and CHOP) after 8h of leucine starvation (0.45mM versus 0mM), a classical inducer of the GCN2 pathway, with or without GCN2i at the indicated concentrations. Tubulin served as a loading control.

B. Western blot analysis of ATF4 after 8h of glutamine deprivation (Gln 2mM versus 0.1mM) with or without GCN2i at the indicated time points. Tubulin served as a loading control.

C. RT-qPCR analysis of canonical ATF4 target genes (*TRB3, CHOP, GADD34, ASNS, SESN2*) upon low glutamine in combination with DMSO or GCN2i. *P58IPK* is given as a non-ATF4 target gene.

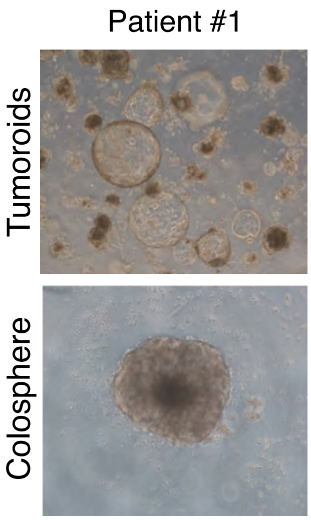

**Figure supplement 4. Illustration of patient-derived tumoroids grown in Matrigel® or as a colosphere in ultra-low attachment plate.**

Microscopy images of the morphology of patient's derived COAD cells grown as matrix-embedded cultures (tumoroids) or in ULA attachment plates (colospheres). Of note, primary cells grown as colosphere lead a visible compaction area.

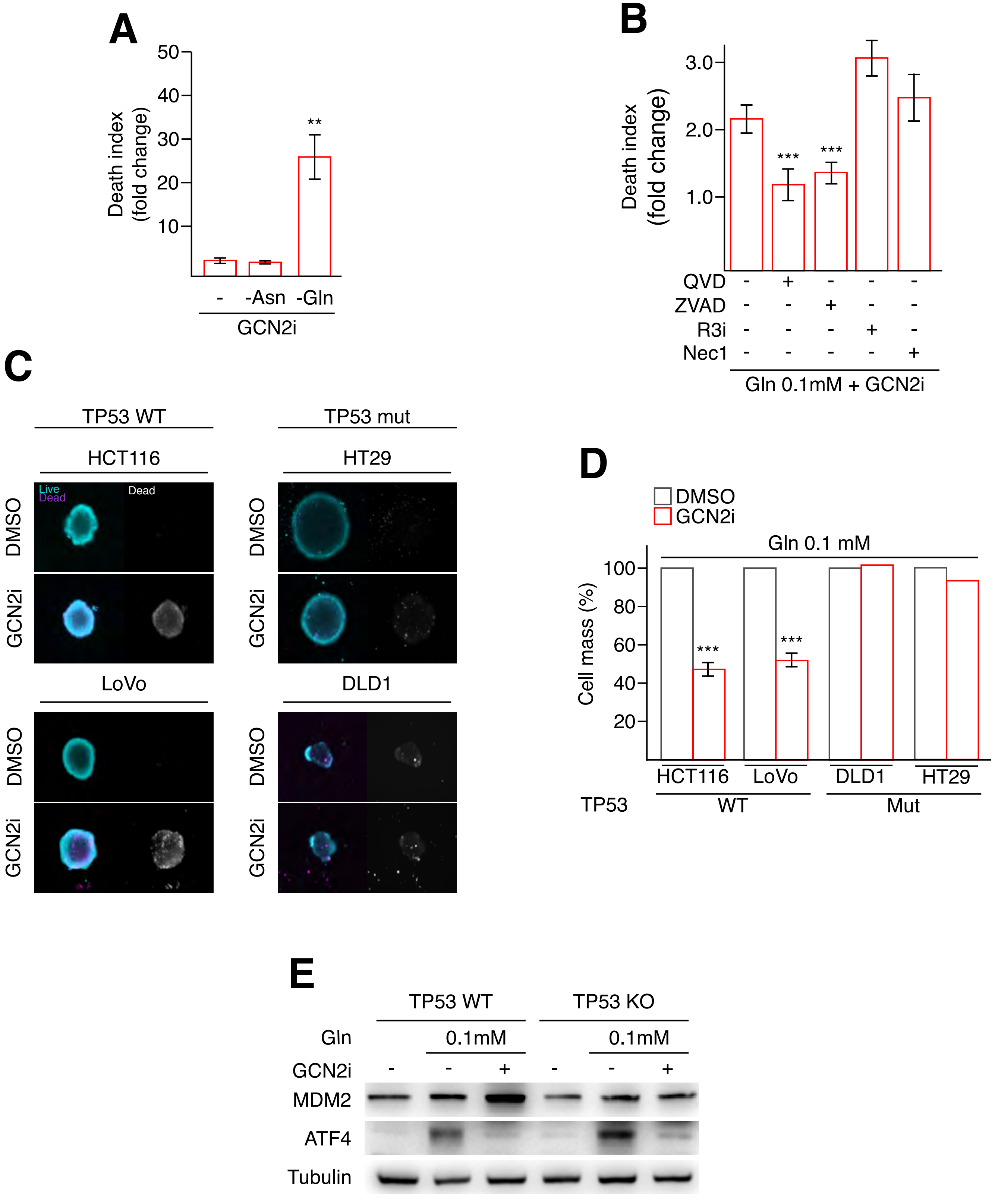

**Figure supplement 5. Lack of GCN2 activity triggers a TP53-dependent cell death upon low glutamine.**

A. HCT116 cell-death analysis following 4 days of GCN2i treatment in complete (-, 2mM Gln, 0.34mM Asn) or starved for glutamine (Gln, 0.1mM) or asparagine (Asn, 0.02mM) media.

B. Impact of inhibitors of necroptosis (Nec1, R3i) or apoptosis (z-VAD, QVD) on GCN2i-induced cell death following a 48h deprivation of glutamine (0.1 mM)

C. Live-and-dead assays in HCT116, LoVo, HT29 and DLD1 spheroids treated for 3 days with vehicle (DMSO) or GCN2i. Enlargment of the spheroids core is provided to visualize cell death.

D. Cell mass measurement of HCT116, LoVo, DLD-1 and HT-29 cells cultured in 2D conditions and treated with DMSO or GCN2i for 48 h. Media were refreshed daily (black rounded arrow).

E. Western blot analysis of ATF4 and MDM2 amounts after 8h of glutamine starvation with or without GCN2i, in HCT116 wild-type (WT) or knockout (KO) for TP53. Tubulin served as a loading control.

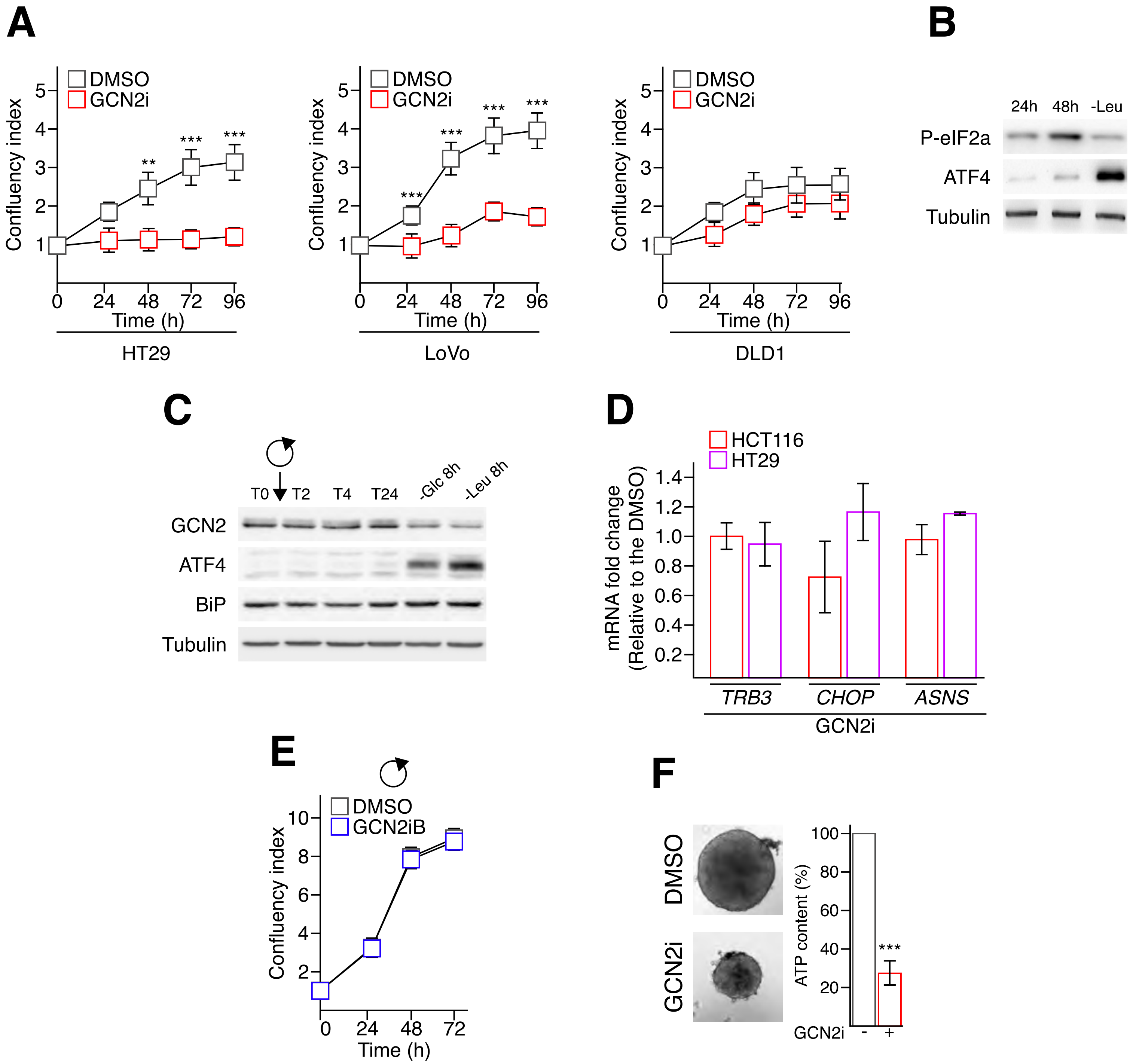

**Figure supplement 6. Lack of GCN2 activity impairs cell proliferation and spheroid formation.**

A. Cell proliferation assessed by the confluency index of HT29, LoVo and DLD1 cells treated or not with GCN2i for 96 hours.

B. Western blot analysis of P-eIF2a and ATF4 amounts after 24 and 48h of culture of HCT116 cells. HCT116 deprived for leucine (Leu) for 8h were used as positive control. Tubulin served as a loading control.

C. Time course analysis of GCN2, ATF4 and BiP amounts in HCT116 cells. HCT116 deprived for leucine (Leu) or glucose (Glc) for 8h were used as positive controls. Tubulin served as a loading control.

D. RT-qPCR analysis of ATF4 target genes (*TRB3, CHOP* and *ASNS*) in HCT116 cells grown in complete medium and treated with DMSO or GCN2i for 48h.

E. Cell proliferation measurement was assessed by the confluency index. HCT116 were treated or not with GCN2iB for 72 hours and media were refreshed every 24 hours.

F. Spheroid formation assays of HT29 treated with DMSO or GCN2i. ATP content was measured after 7 days of treatment. Treatment was initiated concomitantly to plating.

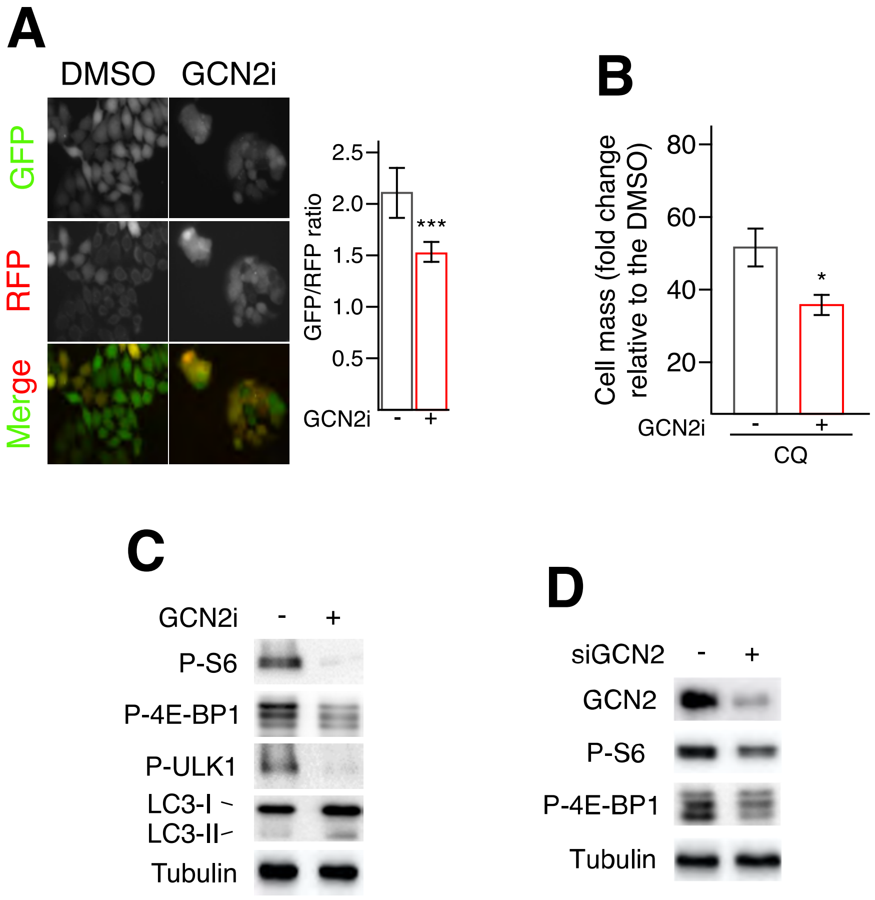

**Figure supplement 7. Lack of GCN2 activity induces an autophagic flux and mTORC1 inhibition in HT29 cells grown in nutrient-rich conditions.**

A. Autophagic flux analysis using HT29 cells stably expressing the GFP-LC3-RFP-LC3ΔG construct and treated with GCN2i for 48 h. Cleavage of GFP-LC3 by autophagy released the RFP-LC3 as an internal control, thus reduction of the GFP/RFP ratio illustrates induction of the autophagic flux.

B. Cell mass was measured in GCN2i-treated HT29 cells after 48 h in combination with chloroquine. Data are expressed relative to the vehicle.

C. Western blot analysis of mTORC1 and autophagy markers (P-S6, P-4E-BP1, P-ULK1, LC3-I/II) following GCN2i treatment for 24h. Tubulin served as a loading control.

D. Western blot analysis of mTORC1 markers (P-S6, P-4E-BP1) following GCN2 silencing for 96h. Tubulin served as a loading control.

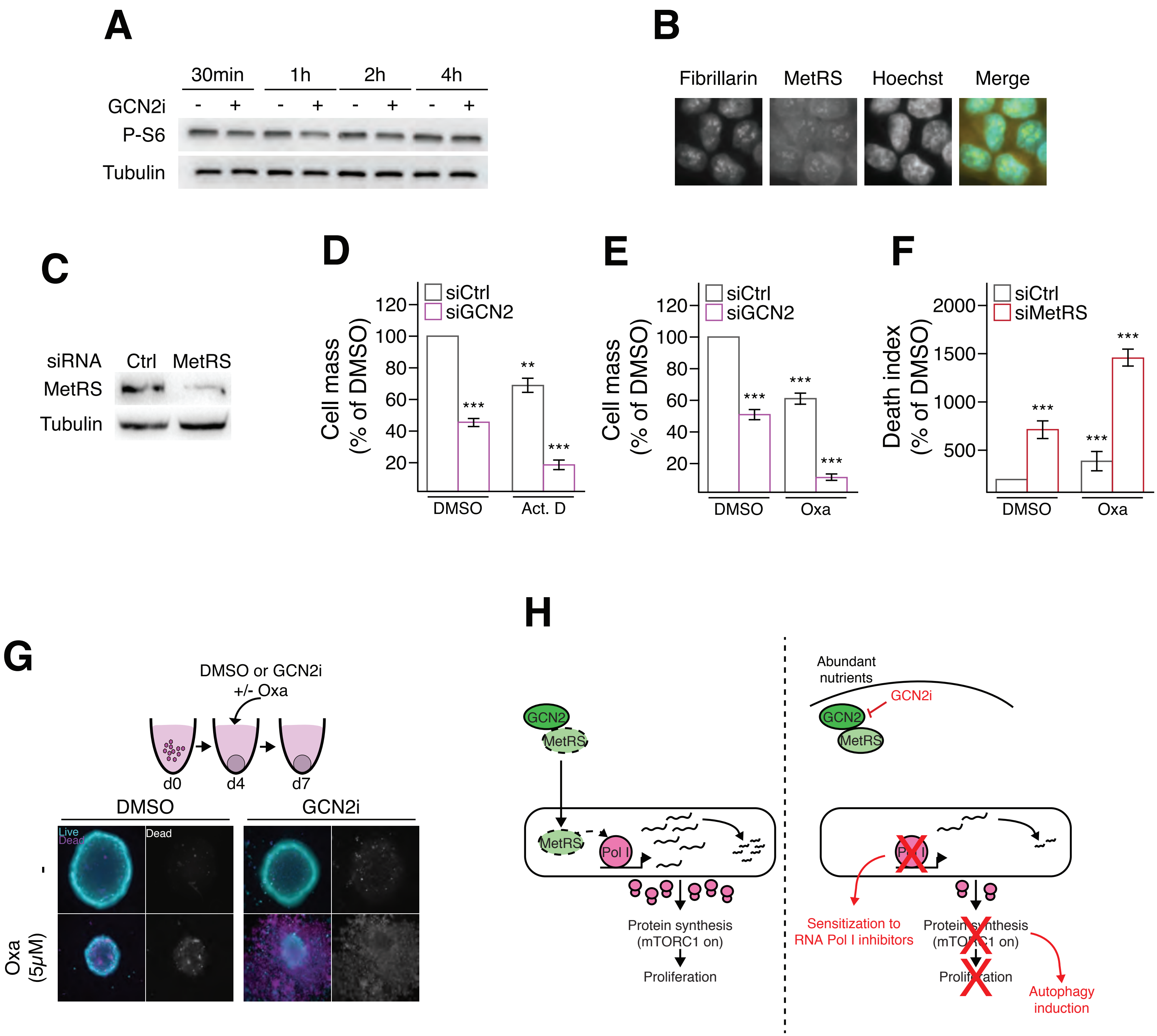

**Figure supplement 8. Lack of GCN2 activity sensitizes COAD cells to oxaliplatin.**

A. Western blot analysis of P-S6 in HCT116 cells treated with or without GCN2i at the indicated time points.

B. Immunocytofluorescence in HCT116 against fibrillarin and methionyl-tRNA synthetase (MetRS) associated to Hoechst coloration.

C. Western blot analysis of MetRS amount in HCT116 cells transfected for 72h with a siRNA control (Ctrl) or against MetRS.

D. Cell mass measurement of HCT116 silenced or not for GCN2 and treated for 48 h with actinomycin D (Act. D).

E. Cell mass measurement of HCT116 silenced or not for GCN2 and treated for 48 h with oxaliplatin (Oxa).

F. Cell death measurement of HCT116 silenced or not for MetRS and treated for 48 h with oxaliplatin (Oxa).

G. Live-and-dead assays in HCT116 spheroids after 5 days of treatment with oxaliplatin (Oxa) and/or GCN2i. To facilitate the observation of cell death induction, staining of non-viable cells is also represented in shades of gray.

H. Schematic model illustrating the role of GCN2 in the control of 47S synthesis in proliferative cells.

| COAD ribosome dysfunction | Amino acid deprivation | COAD Hypoxia | ATF4 |
|---|---|---|---|
| *Negatively-regulated genes* | ASNS | BCCIP | DDIT4 |
| RPL34 | ASS1 | BNIP3L | NARS1 |
| RPL9 | ATF3 | GADD45B | PSAT1 |
| RPL21 | ATF5 | INSIG2 | SREBF1 |
| RPS25 | CARS1 | MPHOSPH6 | IARS1 |
| RPL3 | CBS | TP53 | IARS2 |
| RPL11 | CCNG2 | | MTHFD2 |
| RPL26 | CDKN1A | | EIF2AK4 |
| RPL3L | CEBPB | | ATF3 |
| RPS4Y2 | CHAC1 | | GFPT1 |
| RPL15 | CLEC7A | | WARS |
| RPL22 | CTH | | WARS2 |
| RPS4Y1 | CXCL8 | | NFE2L2 |
| RPL10L | DDIT3 | | YARS |
| RPS27L | DDIT4 | | YARS2 |
| RPL36AL | FYN | | |
| FAU | GADD45A | | |
| RPS29 | MARS | | |
| RPS26 | PPP1R15A | | |
| RPL15 | PSAT1 | | |
| RPS12 | RETN | | |
| RPL10A | SARS | | |
| RPS27A | SESN2 | | |
| RPS27 | SLC38A2 | | |
| RPS13 | SLC7A11 | | |
| RPS23 | STC2 | | |
| RPS3A | TRIB3 | | |
| RPL17 | VEGFA | | |
| RPL39L | WARS | | |
| *Positively-regulated genes* | | | |
| RPL36A | | | |
| RPS2 | | | |
| RPL22L1 | | | |
| RPL8 | | | |
| RPL28 | | | |
| RPLP0 | | | |
| RPS19 | | | |

**Supplementary table 1. List of gene signatures used in the present study.**

| | |
|---|---|
| *ALDH1A3* | Fw: CTGCTACAACGCCCTCTATGCA<br>Rev: GTCGCCAAGTTTGATGGTGACAG |
| *c-MYC* | Fw: CCTGGTGCTCCATGAGGAGAC<br>Rev: CAGACTCTGACCTTTTGCCAGG |
| *TRB3* | Fw: TGGTACCCAGCTCCTCTACG<br>Rev: GACAAAGCGACACAGCTTGA |
| *ASNS* | Fw: CTGTGAAGAACAACCTCAGGATC<br>Rev: AACAGAGTGGCAGCAACCAAGC |
| *P58IPK* | Fw: GCTGAATGTGGAGTAAATGCAG<br>Rev: GTGCAGCTTTTGATTTGCCC |
| *SESN2* | Fw: AGATGGAGAGCCGCTTTGAGCT<br>Rev: CCGATGAAGTCCTCATATCCG |
| *CHOP* | Fw: GGTATGAGGACCTGCAAGAGGT<br>Rev: CTTGTGACCTCTGCTGGTTCTG |
| *GADD34* | Fw: CTGTGATCGCTTCTGGCA<br>Rev: GGAAGAAAGGGTGGGCATC |
| *47S* | Fw: GTGCGTGTCAGGCGTTC<br>Rev: GGGAGAGGAGCAGACGAG |
| *RPS11* | Fw: TGACCTTGATTTATTTTGCATACC<br>Rev: CGAGCAAGACGTTCAGTCCT |

**Supplementary table 2. List of human primers used for RT-qPCR experiments.**

| | |
|---|---|
| *Actin* | CGGTAGGACGCAGACCTGGACCGACCGGCCCTGG |
| *5'ETS* | CGACTGTGCGACAGGAGACCGCTGGACAGCAGCCTCTCCAACCCGGAGGC |
| *ITS2* | ACGCGCCGACCCCCAAGGGAGCGTC |

**Supplementary table 3. List of human probes used for Northern blotting.**

**Supplemental movie 1. Live-imaging of cell death caused by the GCN2 inhibitor in combination with oxaliplatin.** Cell death visualization by incorporation of propidium iodide in HCT116 spheroids treated with the vehicule (DMSO) or GCN2 inhibitor (GCN2i) in combination or not with oxaliplatin (time course of treatments: 120 hours).
